# Histone Lactylation Drives CD8 T Cell Metabolism and Function

**DOI:** 10.1101/2023.08.25.554830

**Authors:** Deblina Raychaudhuri, Pratishtha Singh, Mercedes Hennessey, Bidisha Chakraborty, Aminah J. Tannir, Abel Trujillo-Ocampo, Jin Seon Im, Sangeeta Goswami

## Abstract

The activation and functional differentiation of CD8 T cells are linked to metabolic pathways that result in the production of lactate. Lactylation is a lactate-derived histone post-translational modification (hPTM); however, the relevance of histone lactylation in the context of CD8 T cell activation and function is not known. Here, we show the enrichment of H3K18-lactylation (H3K18la) and H3K9-lactylation (H3K9la) in human and murine CD8 T cells which act as transcription initiators of key genes regulating CD8 T cell phenotype and function. Further, we note distinct impacts of H3K18la and H3K9la on CD8 T cell subsets linked to their specific metabolic profiles. Importantly, we demonstrate that modulation of H3K18la and H3K9la by targeting metabolic and epigenetic pathways regulates CD8 T cell effector function including anti-tumor immunity in preclinical models. Overall, our study uncovers the unique contributions of H3K18la and H3K9la in modulating CD8 T cell phenotype and function intricately associated with metabolic state.

## Introduction

CD8 T cells are critical for mounting an effective immune response against infections and tumors by eliminating pathogen-infected cells and cancer cells(St Paul and Ohashi, 2020; van der Leun et al., 2020; Wong and Pamer, 2003). A naïve CD8 T cell following recognition of a target cell displaying cognate antigen-MHC-I complex becomes activated and proliferates to generate effector CD8 T cells expressing cytotoxic molecules(Harty et al., 2000). T cells acquire metabolic adaptations to support their proliferation and effector functions. While resting T cells primarily depend on the mitochondrial metabolic pathways including tricarboxylic acid (TCA) cycle and oxidative phosphorylation (OXPHOS) to meet their energy requirements, activated T cells undergo a shift toward glycolysis(Geltink et al., 2018; MacIver et al., 2013). Upregulation of glycolysis in activated CD8 T cells leads to increased generation of lactate(Grist et al., 2018). However, the effect of lactate on the epigenetic landscape and function of CD8 T cells is not known.

Lactylation of histone lysine residues is a newly discovered epigenetic modification(Zhang et al., 2019). However, the relevance of histone lactylation in the context of CD8 T cell activation and function is not understood. In this study, we aimed to investigate the role of histone lactylation in the regulation of CD8 T cells and delineate its functional implications. We demonstrated the enrichment of H3K18-lactylation (H3K18la) and H3K9-lactylation (H3K9la) following activation of human and murine CD8 T cells. We also demonstrated the role of H3K18la and H3K9la in CD8 T cells in the context of diseases characterized by aberrant T cell activation including a preclinical model of acute Graft versus Host Disease (GvHD). Furthermore, we investigated the impact of immune checkpoint inhibitors on H3K18la and H3K9la levels in intratumoral CD8 T cells in an *in vivo* preclinical tumor model. We also uncovered the role of H3K18la and H3K9la as transcription regulators in CD8 T cells, individually and in combination with other well-established active histone post-translational modifications (hPTMs). Importantly, we integrated chromatin immunoprecipitation sequencing (ChIP-seq) and RNA-sequencing (RNA-seq) data to show that H3K18la and H3K9la are the primary regulators (in comparison to all other histone PTMs studied) of several genes governing T cell phenotype and function including *Stat1, Cd28*, *Tcf7, Ccr7* and *Batf3*. Importantly, we noted that while H3K18la marks genes critical for activated CD8 T cells, H3K9la is enriched across naïve, activated, and memory subsets. However, the pattern of genes marked by H3K9la varies in different CD8 T cell subsets. Of note, minimal enrichment of H3K18la and H3K9la marks was noted in terminally exhausted CD8 T cells.

Additionally, our data demonstrated that H3K9la and H3K18la are linked to the metabolic status of different CD8 T cell subsets. Pharmacological inhibition of individual cellular metabolic pathways demonstrated that glycolysis primarily drives both H3K18la and H3K9la in activated CD8 T cells while mitochondrial metabolic pathways selectively drive H3K9la in memory CD8 T cells. Further, our data showed that H3K9la is associated with mitochondrial fusion while H3K18la is linked with mitochondrial fission that contributes to the distinct energy metabolism pathways observed in different T cell states.

Furthermore, inhibition of H3K18la and H3K9la by targeting metabolic and epigenetic pathways resulted in decreased expression of T cell effector genes with concurrent inhibition of CD8 T cell-mediated cytotoxicity. Importantly, inhibition of histone delactylases resulted in the enrichment of H3K18la and H3K9la in intratumoral CD8 T cells with concomitant attenuation of tumor growth and increased infiltration of effector CD8 T cells. These findings highlighted the critical role of histone lactylation in regulating CD8 T cell mediated anti-tumor immunity *in vivo*.

Overall, this study uncovers a fundamental role of histone lactylation in regulating the epigenetic, and transcriptomic profiles of different CD8 T cell subsets and provides insight into the distinct roles of H3K18la and H3K9la in modulating CD8 T cell phenotype and function intricately linked to their metabolic state.

## Results

### T cell activation drives histone lactylation

To investigate the relevance of H3K18la and H3K9la in human peripheral CD8 T cells, we assessed the pattern of histone lactylation in naïve and effector CD8 T cells. We stimulated human CD8 T cells isolated from peripheral blood mononuclear cells (PBMCs) with anti-CD3 plus anti-CD28. Flow cytometry analysis revealed an increase in H3K18la and H3K9la in CD8 T cells in activated CD8 T cells compared to naïve CD8 T cells (Figure 1A). Notably, Gzmb+CD69+ activated CD8 T cells exhibited higher levels of H3K18la and H3K9la compared to Gzmb-CD69-CD8 T cells (Figure S1A) supporting a direct association between T cell activation and histone lactylation. Similarly, anti-CD3 plus anti-CD28 mediated activation of murine splenocytes enhanced H3K18la and H3K9la in CD8 T cells (Figure 1B, Figure S1B), mirroring the findings from the human study. Of note, we measured intracellular and extracellular L-lactate levels following CD8 T cell activation. Our findings revealed a time-dependent accumulation of intracellular lactate and extracellular lactate which correlated with the pattern of H3K18la and H3K9la enrichment subsequent to anti-CD3 plus anti-CD28 stimulation (Figure S1C,D, Figure 1B).

**Figure 1:**
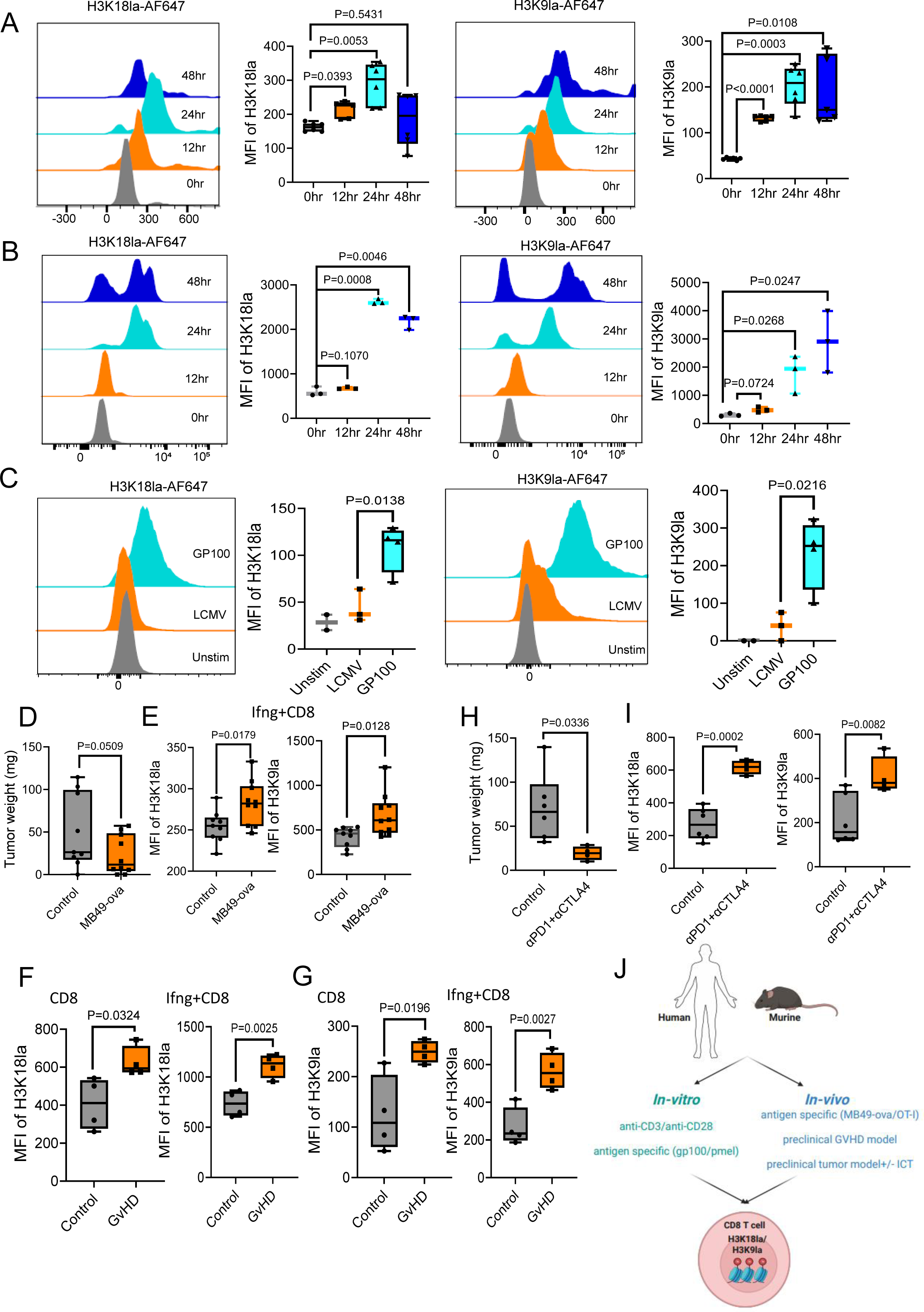
T cell activation drives histone lactylation. (**A**) Representative histograms and box- and-whisker plots demonstrating H3K18la and H3K9la levels (median fluorescence intensity-MFI) in human CD8 T cells following indicated hours of anti-CD3 plus anti-CD28 stimulation. n=5 human donors. Two-tailed Student’s t-test was performed. (**B**) Representative histograms and box-and-whisker plots showing H3K18la and H3K9la levels in murine CD8 T cells following indicated hours of CD3/CD28 stimulation. n= 3 female mice. Two-tailed Student’s t-test was performed. (**C**) Representative histograms and box-and-whisker plots depicting H3K18la and H3K9la levels in pmel mice-derived CD8 T cells stimulated with the cognate GP100 peptide or the control LCMV peptide. n= 3 female mice. Two-tailed Student’s t-test was performed. (**D**) Box- and-whisker plots demonstrating the weights of MB49 tumors and MB49-ova tumors derived from OT-I mice. n=9 female mice bearing control MB49 tumors and 8 female mice bearing MB49-ova tumors. One-tailed Student’s t-test was performed. (**E**) Box-and-whisker plots demonstrating H3K18la and H3K9la levels in TDLN-derived Ifng+ CD8 T cells from MB49 and MB49-ova tumor-bearing mice. n=9 female mice bearing MB49 tumors and 8 female mice bearing MB49-ova tumors. Two-tailed Student’s t-test was performed. (**F**) Box-and-whisker plots demonstrating the level of H3K18la in all CD8 T cells and Ifng+ CD8 T cells, derived from the livers of control and GvHD mice. n= 4 control female mice and 4 GvHD female mice. Two-tailed Student’s t-test was performed. (**G**) Box-and-whisker plots demonstrating the level of H3K9la in all CD8 T cells and Ifng+ CD8 T cells, derived from the livers of control and GvHD mice. n= 4 control female mice and 4 female GvHD mice. Two-tailed Student’s t-test was performed. (**H**) Box-and-whisker plots displaying the weights of MB49 tumors derived from control C57BL/6 mice and mice treated with a combination of anti-PD-1 and anti-CTLA4. n= 6 untreated female mice and 4 combination therapy treated mice. Two-tailed Student’s t-test was performed. (**I**) Box-and-whisker plots depicting H3K18la and H3K9la levels in MB49 tumor-derived CD8 T cells from control and combination therapy treated mice. n= 6 untreated female mice and 4 combination therapy treated mice. Two-tailed Student’s t-test was performed. (**J**) Schematic outlining the *in vitro* and *in vivo* model systems utilized in this study to probe the alterations in H3K18la and H3K9la following CD8 T cell activation. Schematic was created with BioRender.com. Whiskers for all box-and-whisker plots in Figure 1 represent minimum-maximum values. (See also Figure S1, S2, S3)

Further, to interrogate the impact of antigen-specific CD8 T cell stimulation on H3K18la and H3K9la we used CD8 T cells isolated from pmel-1 mice, a model widely used for studying antigen-specific T cell responses(Overwijk et al., 2003). Upon stimulation with the cognate GP100 antigen we observed a robust increase in H3K18la and H3K9la levels in pmel CD8 T cells especially in Gzmb+Tbet+ CD8 T cells while stimulation with the non-cognate lymphocytic choriomeningitis virus (LCMV) peptide failed to induce significant alterations in H3K18la and H3K9la which demonstrated the antigen-specificity of TCR signaling driven changes in H3K18la and H3K9la in CD8 T cells (Figure 1C, Figure S1E). Next, to establish a causal relationship between T cell activation and H3K18la and H3K9la we used a murine *Bhlhe40* knockout model (Bhlhe40 KO) that harbors defects in T cell activation(Li et al., 2019; Salmon et al., 2022). Notably, compared to controls, *Bhlhe40* KO CD8 T cells exhibited significantly reduced levels of H3K18la and H3K9la following anti-CD3 plus anti-CD28 stimulation (Figure S1F) highlighting the impact of impaired T cell activation on H3K18la and H3K9la in CD8 T cells.

Next, to determine the *in vivo* relevance of H3K18la and H3K9la in CD8 T cells we used the OT-I murine model which harbors CD8 T cells bearing T cell receptors (TCRs) specific for the ova-antigen(Hogquist et al., 1994). We injected MB49 tumor cells engineered to express the ova peptide (MB49-ova) into the OT-I mice. We noted attenuated tumor growth of MB49-ova tumors compared to control MB49 tumors (Figure 1D). Further, analyses of flow cytometry data demonstrated higher H3K18la and H3K9la levels in Ifng+ CD8 T cells within the tumor-draining lymph nodes (TDLNs) of mice bearing MB49-ova tumors compared to control MB49 tumors (Figure 1E, Figure S1G, H). Next, we interrogated the relevance of H3K18la and H3K9la in preclinical disease models including acute graft versus host disease (GvHD) and a tumor model following treatment with immune checkpoint therapy. The hyperactivation of CD8 T cells is a key driver of GvHD where these overactive T cells attack the host’s tissues and organs (Jiang et al., 2021; Khandelwal et al., 2020; Lee et al., 2023; Zheng et al., 2009). On the other hand, CD8 T cell activation is essential for anti-tumor immunity and response to immune-based therapies(Giles et al., 2023).

To establish the GvHD model we conducted bone marrow and splenocyte transplantation from C57BL/6 mice into irradiated BALB/c mice following a previously established protocol(Pareek et al., 2022). 8 days post-transplantation we collected livers to examine organ specific GvHD immune responses in control and GvHD mice. Our primary objective was to evaluate alterations in H3K18la and H3K9la within hyperactivated CD8 T cells originating from GvHD mice. As expected, we observed increased abundance of activated Gzmb+ and Tbet+ CD8 T cells in the livers of GvHD mice indicating heightened inflammation in these animals (Figure S2A-C). We also noted elevated levels of H3K18la and H3K9la in activated Gzmb+ and Ifng+ CD8 T cells compared to their Gzmb- or Ifng-counterparts (Figure S2A, D, E). This observation aligns with our previous *in vitro* findings linking T cell activation with histone lactylation. Importantly, we found significantly higher levels of H3K18la and H3K9la in CD8 T cells as well as in Gzmb+, IFNg+, and Tbet+ activated CD8 T cells derived from the livers of GvHD mice compared to control mice further reinforcing the connection between CD8 T cell activation and histone lactylation in the disease context (Figure 1F, G, Figure S2A, F, G).

Next, we evaluated the H3K18la and H3K9la levels in CD8 T cells within the preclinical MB49 tumor model. Notably, intratumoral Ifng+ activated CD8 T cells harbored higher levels of H3K18la and H3K9la compared to Ifng-CD8 T cells (Figure S3A, B), mirroring both our *in vitro* data and the findings from the GvHD model. We and others have previously demonstrated the sensitivity of MB49 tumors to immune checkpoint inhibitors. Therefore, we evaluated the impact of immune checkpoint therapy, a potent inducer of T cell mediated anti-tumor responses(Sharma et al., 2023; Sharma et al., 2021) on CD8 T cell H3K18la and H3K9la levels in the MB49 tumor model. Similar to previous data we noted reduction in tumor sizes (Figure 1H) and increase in interferon gamma expression in intratumoral CD8 T cells in MB49 tumor-bearing mice treated with a combination of anti-PD-1 and anti-CTLA-4 compared to the control-MB49 tumor-bearing mice (Figure S3C). Importantly, we noted a substantial enhancement of H3K18la and H3K9la in intratumoral CD8 T cells from mice treated with the anti-PD-1 plus anti-CTLA-4 therapy (Figure 1I, Figure S3D) which underscores the critical link between T cell activation and the dynamic regulation of histone lactylation.

Cumulatively, using *in vitro* anti-CD3 plus anti-CD28 stimulation of human and murine CD8 T cells, *in vitro* and *in vivo* antigen-specific stimulation of CD8 T cells derived from pmel and OT-I mice, as well as preclinical disease models including acute graft versus host disease (GvHD) and a preclinical tumor model treated with immune checkpoint therapy (Figure 1J) we demonstrate that CD8 T cell activation which results in the generation of lactate in CD8 T cells, results in the increase of H3K18la and H3K9la.

### Histone lactylation(s) regulate transcription initiation in CD8 T cells

To elucidate the role of H3K18la and H3K9la in the transcriptional regulation of CD8 T cells we performed ChIP-seq using naïve and activated CD8 T cells. ChromHMM analysis(Ernst and Kellis, 2017) of the ChIP-seq data allowed us to classify the genomic regions into 7 distinct chromatin states based on specific combinations of histone modifications including histone H3 lysine 4 trimethylation (H3K4me3), histone H3 lysine 9 acetylation (H3K9ac), histone H3 lysine 27 acetylation (H3K27ac), histone H3 lysine 79 dimethylation (H3K79me2), histone H3 lysine 36 trimethylation (H3K36me3), histone H3 lysine 27 trimethylation (H3K27me3), H3K9la, H3K18la and RNA polymerase II (RNApol II) (Figure 2 A-C). H3K4me3 and H3K9ac are well-established epigenetic marks associated with transcription initiation and H3K27ac marks active enhancer regions(Millan-Zambrano et al., 2022; Talbert and Henikoff, 2021). H3K79me2 and H3K36me3 are associated with actively transcribed gene bodies while H3K27me3 is a mark of transcription repression(Millan-Zambrano et al., 2022; Talbert and Henikoff, 2021). We found that chromatin regions enriched with H3K18la or H3K9la marks display a pattern of active transcription-associated chromatin state characterized by the enrichment of other transcription initiation marks including H3K4me3, H3K9ac, and H3K27ac (Figure 2A). Further, both H3K18la and H3K9la marks exhibit significant enrichment in chromatin regions proximal to transcription start sites (TSS) and CpG islands indicating their involvement in the regulation of transcription initiation and gene expression (Figure 2A-C). Next, to delineate the relationship of H3K18la and H3K9la with the other hPTMs in CD8 T cells, we investigated the correlation of H3K18la and H3K9la with these hPTMs. The correlation matrices revealed a distinct pattern of positive correlations between lactylation marks (H3K18la and H3K9la) and other hPTMs associated with active transcription initiation including H3K4me3 and H3K9ac in promoter regions (Figure 2D). Additionally, lactylation marks showed positive correlations with H3K27ac which marks active enhancer and promoter regions (Figure 2D). We also observed positive correlations between lactylation marks and H3K79me2 and H3K36me3, which mark actively transcribed gene bodies (Figure 2D). Importantly, we noted similar correlations in proximal and distal enhancer-like sequence bearing regions (pELS and dELS) but not in intergenic regions (Figure S4A-C). These findings indicate a potential role of histone lactylation marks in promoting active gene expression and transcriptional regulation in CD8 T cells.

**Figure 2:**
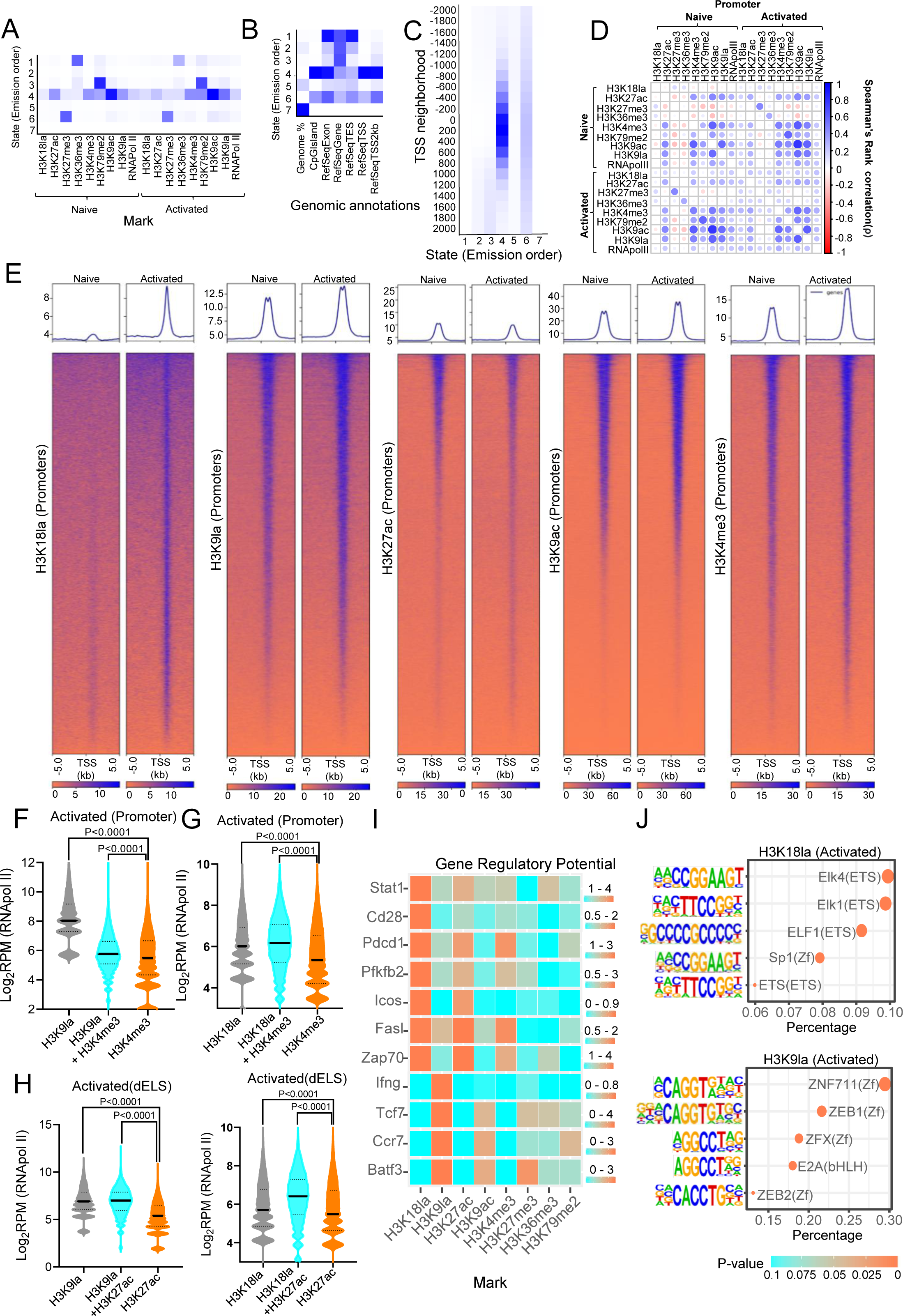
Histone lactylation regulates transcription initiation in CD8 T cells. **(A)** Heatmap depicting emission parameters, generated from ChromHMM analysis. Each row corresponds to a specific chromatin state and each column corresponds to a specific epigenetic mark in naive and activated murine CD8 T cells. **(B)** Heatmap displaying the fold enrichment of different chromatin states in distinct genomic regions based on the indicated epigenetic marks in naïve and activated CD8 T cells. **(C)** Heatmap demonstrating the fold enrichment of different chromatin states around transcription start sites (TSS) based on epigenetic marks shown in **A,** in naïve and activated CD8 T cells. The darker blue color corresponds to a greater probability of observing the mark in the state. **(D)** Correlation matrix demonstrating Spearman’s correlation coefficients (color intensities and the size of circles) among H3K18la and H3K9la sites and the other indicated epigenetic modifications at the promoter regions (TSS±1 kb). **(E)** Heatmaps demonstrating the genomic occupancy of the indicated hPTMs at gene promoter regions in murine naïve and activated CD8 T cells. **(F-H)** Violin plots depicting normalized RNApol II binding (log2RPM) in promoter **(F,G)** and distal enhancer regions of activated CD8 T cells **(H).** Regions have been categorized based on the combination of indicated histone PTMs. Two-tailed Student’s t-test was performed. (**I**) Heatmaps demonstrating the gene regulatory potential of each of the hPTMs for the indicated genes as calculated using BETA. **(J)** Dotplots demonstrating the top 5 significantly enriched transcription factor binding motifs in H3K18la (top panel) and H3K9la (bottom panel) marked promoter regions of activated CD8 T cells. (See also Figure S4, S5)

Next, to gain a more comprehensive understanding of the dynamic alterations of histone modifications during the transition from the naive to activated state of CD8 T cells, we compared the hPTM enrichment in the promoter, proximal enhancer, and distal enhancer regions between naïve and activated CD8 T cells. We noted enrichment of H3K9la, H3K4me3 and H3K9ac in the promoter regions in the naïve T cell state which increased following T cell activation along with increased RNApol II occupancy in the promoter regions (Figure 2E, Figure S4D). Importantly, H3K18la which showed minimal enrichment in the naïve state demonstrated substantial enrichment in the promoter regions of genes following T cell activation (Figure 2E). While H3K27ac is traditionally linked to active enhancers we also noted enrichment of H3K18la and H3K9la marks in proximal enhancer regions (pELS) of genes in activated CD8 T cells, consistent with RNApol II binding (Figure S5A-C, F). This indicates that H3K18la and H3K9la mark both promoter and enhancer regions. Notably, we observed a significant enrichment of H3K18la compared to other hPTMs in the distal enhancer region (Figure S5A, B, D, E) with concurrent increase in RNApol II occupancy in the activated CD8 T cells (Figure S5F). This highlights a critical role of H3K18la in promoting gene transcription through proximal as well as distal enhancer regions.

To determine the implications of promoter and enhancer regions marked by H3K18la or H3K9la we compared RNApol II occupancy of regions marked by unique lactylation peaks, regions marked by unique peaks of other active hPTMs peaks including H3K4me3 and H3K27ac and regions marked by overlapping lactylation peaks and H3K4me3 or H3K27ac peaks. In activated CD8 T cells we observed that gene promoters enriched with unique H3K9la peaks displayed notably higher RNApol II occupancy compared to promoter regions marked solely by H3K4me3 peaks (Figure 2F). Similarly, promoters enriched with unique H3K18la peaks exhibited significantly elevated RNApol II binding in comparison to promoter regions exclusively marked by H3K4me3 peaks (Figure 2G). These findings suggest that in CD8 T cells RNApol II binding to gene promoters carrying H3K9la and H3K18la marks is higher compared to promoter regions marked by H3K4me3 alone. Additionally, distal enhancer-like sequences bearing overlapping H3K9la and H3K27ac peaks or H3K18la and H3K27ac peaks demonstrated higher RNApol II occupancy compared enhancer regions marked solely with H3K27ac peaks (Figure 2H). This suggests a synergistic effect of H3K9la and H3K18la with H3K27ac in driving gene transcription through distal enhancers during CD8 T cell activation.

Next, we assessed the relative contribution of the various hPTMs in governing the transcriptomic profile of activated CD8 T cells. We employed Binding and Expression Target Analysis (BETA)-based integration of ChIP-seq and RNA-seq data to determine the gene expression regulatory potential score associated with each of these hPTMs. The data revealed that H3K18la is the primary regulator of genes crucial for T cell activation including *Stat1, Cd28, Icos, Pdcd1* and *Pfkfb2* (Figure 2I). H3K18la also regulates other T cell activation genes including *Fasl* and *Zap70* in conjunction with H3K27ac and H3K4me3 (Figure 2I). On the other hand, H3K9la is the primary regulator of the T cell activation gene *Ifng* as well as for genes associated with naïve and memory T cells including *Tcf7, Ccr7* and *Batf3* (Figure 2I). Overall, this analysis demonstrated the importance of H3K18la and H3K9la as the major regulators of several genes critical for governing T cell phenotype and function.

To further investigate the interactions of transcription factors (TFs) with H3K18la and H3K9la marked promoter regions, we conducted a comprehensive TF binding motif enrichment analysis using Hypergeometric Optimization of Motif EnRichment (HOMER)(Heinz et al., 2010). Investigation of the top 5 most enriched TF binding motifs demonstrated that H3K18la marked promoter regions are enriched in ETS transcription factor binding motifs while H3K9la marked promoter regions are enriched for zinc finger (ZF) transcription factor binding motifs, in activated CD8 T cells (Figure 2J). It is noteworthy that both the ETS and ZF transcription factors have been extensively recognized for their pivotal roles in shaping CD8 T cell differentiation, homeostasis, and functional outcomes(Guan et al., 2018; Luo et al., 2017; Muthusamy et al., 1995; Schauder et al., 2021; Smith-Raska et al., 2018; Wang et al., 1993).

Cumulatively, the data highlights a critical role of H3K18la and H3K9la as the primary regulators of several key genes governing T cell phenotype and function. The data also reveal a distinct pattern of TF binding motif enrichment in H3K18la and H3K9la marked promoter regions. Overall, our findings demonstrate a complex interplay of H3K18la and H3K9la marks with other hPTMs and TFs in driving transcription in CD8 T cells.

### Distinct patterns of H3K9la and H3K18la enrichment in different CD8 T cell subsets

Based on our data that demonstrated H3K9la enrichment in both naive and activated states while H3K18la enrichment was specifically noted in activated CD8 T cells (Figure 2E), we investigated the differential enrichment of H3K18la and H3K9la across specific gene loci within naïve and activated states. In naive CD8 T cells, we observed enrichment of H3K9la in multiple genes involved in maintaining T cell quiescence such as *Bach2, Foxo1, Klf2* whereas in activated CD8 T cells effector genes including *Gzmk, Gzmb, Prf1, Icos, Fasl,* and *Pdcd1* displayed distinct enrichment of H3K18la marks (Figure 3A, Figure S6A). Next, to delineate the enrichment of H3K18la and H3K9la in memory CD8 T cells, we generated memory CD8 T cells through prolonged IL-15 stimulation of activated T cells(Pilipow et al., 2015; Richer et al., 2015). Comparative assessment of H3K18la and H3K9la enrichment in memory CD8 T cells revealed selective enrichment of H3K9la peaks in genes associated with memory T cells including *Lef1, Ccr7,* and *Il7r*, similar to the pattern observed in naive T cells (Figure 3B, Figure S6A).

**Figure 3:**
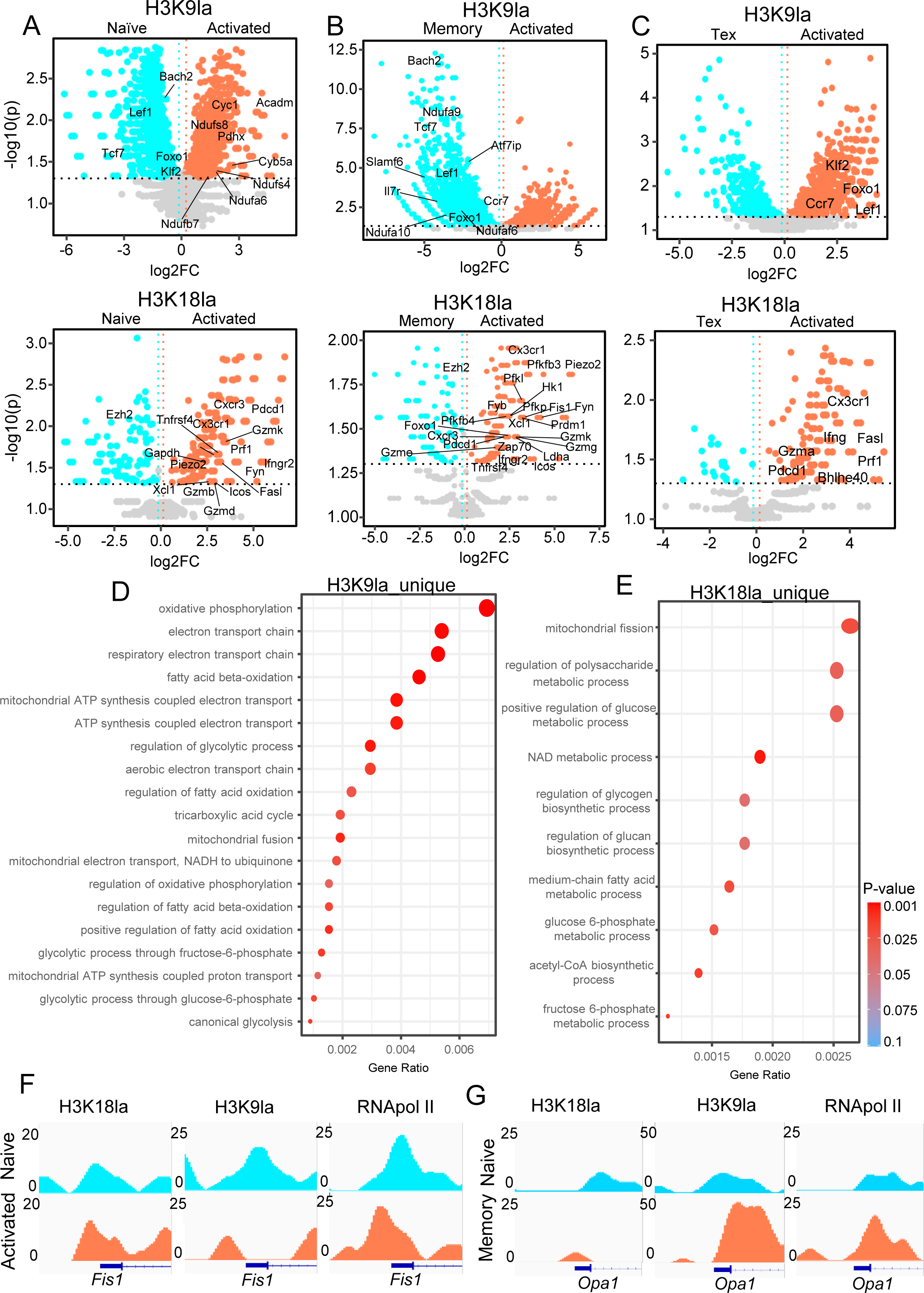
Distinct patterns of H3K9la and H3K18la enrichment in different CD8 T cell subsets. **(A-C)** Volcano plots representing genes harboring differentially enriched H3K9la (upper panel) or H3K18la marks (lower panel) in naïve versus activated **(A)**, memory versus activated **(B),** and terminally exhausted (Tex) versus activated **(C)** CD8 T cells. Volcano plots show the log2 ratio of fold change (log2FC) plotted against the Absolute Confidence, -log10 adjusted p-value (-log10(p)). The p-value was derived from the Student’s t-test using MAnorm. **(D,E)** Dotplots representing cellular metabolic pathways obtained from GO enrichment analysis on gene sets bearing unique H3K9la peaks in naïve CD8 T cells **(D)** and unique H3K18la peaks in activated CD8 T cells **(E).** The p-values were calculated by hypergeometric test using ChIPpeakAnno. **(F,G)** Genome Browser view of H3K18la, H3K9la, and RNApol II peaks at the *Fis1* **(F)** and *Opa1* **(G)** gene promoter regions (TSS ± 1kb) in the indicated CD8 T cell subsets. (See also Figure S6)

Additionally, to interrogate the role of H3K18la and H3K9la in exhausted CD8 T cells, we generated terminally exhausted CD8 T cells (Tex) *in vitro(Belk et al., 2022)*. We noted depletion of H3K18la and H3K9la marks in exhausted CD8 T cells compared to activated and memory subsets (Figure 3C, Figure S6B). Further, we did not note enrichment of H3K18la and H3K9la marks in exhaustion-associated genes including *Lag3* and *Entpd1* (Figure S6C). To confirm the H3K18la and H3K9la levels in exhausted CD8 T cells *in vivo*, we assessed the levels of H3K18la and H3K9la levels in exhausted intratumoral CD8 T cells from MB49 tumor-bearing mice. We noted lower levels of H3K18la and H3K9la in Lag3+ exhausted intratumoral CD8 T cells compared to Tbet+ effector CD8 T cells mirroring the histone lactylation pattern observed in *in vitro* generated exhausted CD8 T cells (Figure S6D). Further analysis of other hPTMs revealed enrichment of H3K9ac in exhaustion-associated genes in exhausted CD8 T cells generated *in vitro* (Figure S6C, E).

To investigate the distinct patterns of H3K18la and H3K9la enrichment across key genetic loci in naïve, activated, and memory CD8 T cells, we performed Gene Set Enrichment Analyses (GSEA)(Subramanian et al., 2005) of the differentially enriched H3K18la and H3K9la marked gene loci between these CD8 T cell states. Notably, H3K18la marks correlated with heightened inflammatory response and Jak-Stat signaling pathways upon T cell activation (Figure S6F). H3K9la marks were associated with upregulated oxidative phosphorylation pathway along with downregulated TNFα signaling and cytokine-cytokine receptor interaction pathways in activated T cells compared to their naive counterparts (Figure S6G). GSEA also linked H3K9la marks to upregulated inflammatory response and fatty acid metabolism pathways in memory CD8 T cells (Figure S6H).

Importantly, our analysis revealed that in activated CD8 T cells H3K9la marks are enriched in genes involved in OXPHOS including *Ndufs8, Ndufa6,* and *Ndufs4* while H3K18la marks are enriched in genes associated with glycolysis including *Hk1, Pfkl, and Pfkp* indicating a role of H3K18la and H3K9la in regulating specific metabolic pathways in activated CD8 T cells (Figure 3A, Figure S6A). Therefore, we interrogated the unique and overlapping peaks of H3K18la and H3K9la marks and annotated the genes specifically associated with these distinct peaks. Consistent with the differential enrichment analysis we found that unique H3K9la peaks are enriched in mitochondrial metabolism pathways including oxidative phosphorylation, electron transport chain, and fatty acid beta oxidation in naive CD8 T cells (Figure 3D). Conversely, unique H3K18la peaks showed enrichment in glucose metabolism pathways in activated CD8 T cells (Figure 3E). These findings demonstrated that H3K9la plays a role in modulating mitochondrial metabolism in CD8 T cells while H3K18la is particularly involved in regulating the glucose metabolism during CD8 T cell activation. Importantly, we observed that H3K18la marks are enriched in the mitochondrial fission pathway in activated CD8 T cells while H3K9la marks show prominence in the mitochondrial fusion pathway in naïve CD8 T cells (Figure 3D, E). Further, we found that the promoter regions of *Fis1*, a gene responsible for mitochondrial fission(Kim et al., 2011; Yu et al., 2019), exhibited enrichment of H3K18la in activated CD8 T cells (Figure 3F). In contrast, *Opa1*, known for promoting mitochondrial fusion in memory CD8 T cells(Buck et al., 2015), displayed significant enrichment of H3K9la corresponding to increased RNApol II levels in the memory state (Figure 3G). This suggests that H3K18la possibly plays a crucial role in regulating mitochondrial fission during CD8 T cell activation while H3K9la plays a pivotal role in governing mitochondrial fusion during the formation of memory CD8 T cells. These findings highlight that H3K18la and H3K9la play an important role in governing mitochondrial dynamics in T cells, including mitochondrial fission and fusion which contribute to the distinct energy metabolism pathways observed in different T cell states(Buck et al., 2015; Rambold and Pearce, 2018; Simula et al., 2018).

Taken together, the data demonstrate that while H3K18la marks genes critical for activated CD8 T cells, H3K9la is enriched across naïve, activated, and memory subsets. However, the pattern of genes marked by H3K9la varies in different CD8 T cell subsets. Further, minimal enrichment of H3K18la and H3K9la marks was noted in terminally exhausted CD8 T cells.

### Histone lactylation is linked to the metabolic status of distinct CD8 T cell subsets

While activated CD8 T cells primarily rely on endogenous glycolysis for ATP generation, naïve and memory subsets depend on mitochondrial metabolic pathways including OXPHOS and fatty acid oxidation (FAO) for energy generation(Geltink et al., 2018; Grist et al., 2018; MacIver et al., 2013). To determine the role of endogenous lactate in the regulation of H3K18la and H3K9la following T cell activation, we targeted multiple metabolic enzymes driving endogenous lactate generation in activated CD8 T cells (Figure 4A). First, we treated activated CD8 T cells with a LDHA inhibitor which resulted in substantial attenuation of cellular glycolysis (Figure S7A), accompanied by diminished intracellular L-lactate levels (Figure S7B) and a concurrent depletion of both H3K18la and H3K9la in activated CD8 T cells (Figure 4B, Figure S7C, D). In addition to pharmacological inhibition of LDHA, shRNA mediated knockdown of *Ldha* also attenuated H3K18la and H3K9la levels in activated CD8 T cells, corroborating these results (Figure S7E). Importantly, LDHA inhibition in activated CD8 T cells depleted H3K18la and RNA pol II from the promoter region of *Fis1* with concomitant alteration in the mitochondrial morphology from fission (fragmented pattern) towards mitochondrial fusion (network/diffused pattern) demonstrated by super resolution and transmission electron microscopy (Figure 4C, D, Figure S7F, G).

**Figure 4:**
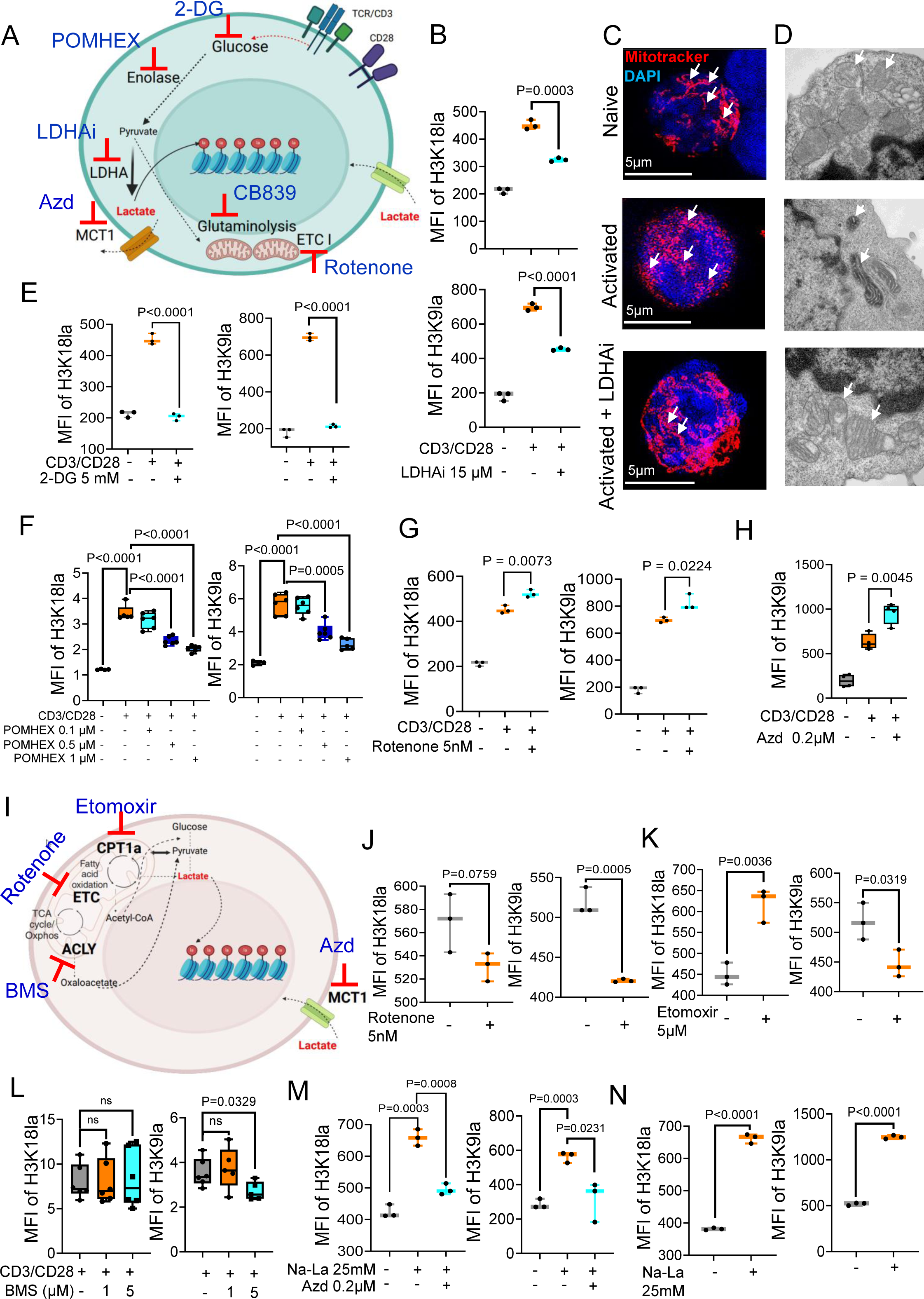
Histone lactylation is linked to the metabolic status of distinct CD8 T cell subsets. **(A)** Schematic representation of various endogenous metabolic pathways contributing to the generation of intracellular lactate and the specific enzymes we targeted to modulate histone lactylation in activated CD8 T cells. Schematic was created with BioRender.com. **(B)** Box-and-whisker plots demonstrating the MFI of H3K18la and H3K9la in naïve, activated and 15 µM LDHA inhibitor treated activated CD8 T cells. (n=3) Two-tailed Student’s t-test was performed. **(C,D)** Representative super-resolution microscopy images **(C)** and transmission electron microscopy images **(D)** of naïve, activated and 15 µM LDHA inhibitor treated activated CD8 T cells depicting mitochondrial morphology (red-MitoTracker) and nuclei (blue-DAPI). Arrows point to fused and fissed mitochondrial morphology. **(E)** Box-and-whisker plots depicting MFI of H3K18la and H3K9la in naïve, activated and 5 mM 2-DG (glucose analog) treated activated murine CD8 T cells. (n=3) Two-tailed Student’s t-test was performed. **(F)** Box-and-whisker plots showing the MFI of H3K18la and H3K9la in naïve, activated, 0.1 µM, 0.5 µM and 1 µM POMHEX (enolase inhibitor) treated activated CD8 T cells. (n=5) Two-tailed Student’s t-test was performed. **(G)** Box-and-whisker plots demonstrating the MFI of H3K18la and H3K9la in naïve, activated and 5 nM Rotenone (ETC-I inhibitor) treated activated CD8 T cells. (n=3) Two-tailed Student’s t-test was performed. **(H)** Box and whisker plot depicting the MFI of H3K9la in naïve, activated and 0.2 µM Azd (MCT1 inhibitor) treated activated CD8 T cells. (n=4) Two-tailed Student’s t-test was performed. **(I)** Schematic representation of various endogenous metabolic pathways and the specific enzymes we inhibited to modulate histone lactylation in memory CD8 T cells. Schematic was created with BioRender.com. **(J)** Box-and-whisker plots demonstrating the MFI of H3K18la and H3K9la in untreated and 5 nM Rotenone (ETC-I inhibitor) treated memory CD8 T cells. (n=3) Two-tailed Student’s t-test was performed. **(K)** Box-and-whisker plots showing the MFI of H3K18la and H3K9la in untreated and 5 µM Etomoxir (Cpt1a inhibitor) treated memory CD8 T cells. (n=3) Two-tailed Student’s t-test was performed. **(L)** Box-and-whisker plots depicting the MFI of H3K18la and H3K9la in activated, 1 µM and 5 µM BMS-303141 (ACLY inhibitor) treated memory CD8 T cells. (n=6) Two-tailed Student’s t-test was performed. ns= non-significant **(M)** Box-and-whisker plots demonstrating the MFI of H3K18la and H3K9la in untreated, 25 mM sodium lactate (Na-La) and 0.2 µM Azd (MCT1 inhibitor) treated naïve CD8 T cells. (n=3) Two-tailed Student’s t-test was performed. **(N)** Box-and-whisker plots demonstrating the MFI of H3K18la and H3K9la in untreated and 25 mM Na-La treated memory CD8 T cells. (n=3) Two-tailed Student’s t-test was performed. Whiskers for all box-and-whisker plots in Figure 4 represent minimum-maximum values. (See also Figure S7, S8)

To further confirm the role of endogenous glycolysis in driving histone lactylation in activated CD8 T cells, we treated CD8 T cells with 2-deoxy glucose (2-DG) and POMHEX during activation (Figure 4A). 2-DG, a glucose analog, attenuates glycolysis by inhibiting hexokinase and glucose-6-phosphate isomerase activities, while POMHEX targets enolase to hinder glycolysis-derived lactate production (Figure S7A, B). Notably, treatment with both 2-DG and POMHEX attenuated H3K18la and H3K9la levels in activated CD8 T cells confirming the role of glycolysis in driving histone lactylation in activated CD8 T cells (Figure 4E, F, Figure S7H). Next, to investigate additional endogenous metabolic pathways contributing to lactate generation in activated CD8 T cells, such as glutaminolysis, we treated CD8 T cells with the Glutaminase 1 inhibitor, CB839 (Figure 4A) which attenuated both H3K18la and H3K9la in activated CD8 T cells (Figure S7I). We also assessed the levels of H3K18la and H3K9la following treatment of activated CD8 T cells with Rotenone, an Electron Transport Chain (ETC-I) inhibitor (Figure 4A) known to induce an increase in intracellular lactate levels(Hou et al., 2018; Karlsson et al., 2016). Importantly treatment with Rotenone led to a significant increase in both H3K18la and H3K9la (Figure 4G, Figure S7J). Additionally, inhibition of monocarboxylate transporter (MCT1) by Azd (Figure 4A) which facilitates lactate transport across cell membranes along their concentration gradients(Brown and Brooks, 1994; Juel, 1996) in activated CD8 T cells increased H3K9la *in vitro* (Figure 4H, Figure S7K) indicating that inhibition of MCT1 prevents export of endogenous lactate, consequently increasing histone lactylation. Thus, by modulating multiple pathways of intracellular lactate generation and by inhibiting lactate transporters (Figure 4A) we confirmed that endogenous metabolic pathways of lactate generation serve as the major source of H3K18la and H3K9la in activated CD8 T cells.

We observed elevated H3K18la levels only in activated CD8 T cells, which are known for their high glycolytic activity. In contrast, the levels of H3K9la remained consistently high in naïve and memory T cells (Figure 3) which do not rely on glycolysis as the primary pathway of energy generation. Therefore, we interrogated the potential differences in the endogenous metabolic sources of H3K9la and H3K18la in memory CD8 T cells. Notably, neither enolase nor LDHA inhibition attenuated histone lactylation in memory CD8 T cells (Figure S8A, B) indicating that glycolysis is not the major driver of H3K18la and H3K9la in the memory subset. Memory CD8 T cells primarily utilize mitochondrial oxidative phosphorylation and FAO for energy generation (Figure 4I)(Geltink et al., 2018; MacIver et al., 2013). Therefore, we also investigated the role of these mitochondrial pathways, including ETC and FAO in governing the levels of H3K18la and H3K9la in memory CD8 T cells (Figure 4I). Unlike in activated CD8 T cells (Figure 4G), inhibition of ETC with Rotenone resulted in a significant decrease selectively in H3K9la levels (Figure 4I, J, Figure S8C). Additionally, inhibition of Cpt1a-mediated FAO using Etomoxir also reduced H3K9la levels, but not H3K18la levels, in memory CD8 T cells (Figure 4I, K, Figure S8D).

ATP citrate lyase (ACLY) is an enzyme which converts mitochondrial citrate to acetyl-CoA and oxaloacetate(Lin et al., 2020; Zheng et al., 2021). Subsequently, malate dehydrogenase and malic enzyme 1 convert oxaloacetate to pyruvate which in turn generates lactate. Therefore, ACLY inhibition, attenuates a pathway of mitochondrial lactate generation (Figure 4I)(Liu et al., 2023). Inhibition of ACLY with BMS-303141 (BMS) caused a selective decrease in H3K9la levels in memory CD8 T cells indicating mitochondrial ACLY-mediated lactate generation as an additional endogenous source of H3K9la in the memory subset (Figure 4L). Together, these findings demonstrate the role of endogenous mitochondrial metabolic pathways in driving H3K9la in memory CD8 T cells. Next, we investigated the impact of H3K9la modulation on the regulation of genes associated with CD8 T cell memory phenotype. We treated CD8 memory T cells with BMS, which attenuated H3K9la enrichment in the promoter regions of genes associated with memory phenotype including *Lef1 and Bach2* with a concomitant reduction in gene expression (Figure S8E, F). Together, these findings show that targeting mitochondrial metabolism pathways selectively regulate H3K9la levels in memory CD8 T cells which in turn regulates the expression of genes associated with the memory phenotype.

Subsequently, to determine the impact of exogenous lactate on H3K18la and H3K9la in CD8 T cells, we supplemented activated CD8 T cell cultures with sodium-L-lactate (Na-La). However, addition of exogenous lactate did not increase H3K18la and H3K9la levels in activated CD8 T cells (Figure S8G, Figure S7C). Contrary to activated CD8 T cells, addition of 25mM exogenous sodium lactate significantly increased H3K18la and H3K9la in both naïve and memory CD8 T cells (Figure 4M, N, Figure S8G). Additionally, MCT1 inhibition attenuated the exogenous lactate induced increase in histone lactylation in naïve CD8 T cells (Figure 4M, Figure S8G). Further, ChIP-seq and RNA-seq analyses of naïve CD8 T cells treated with sodium lactate demonstrated significant enrichment of H3K9la in the promoter regions of several genes associated with the naïve phenotype including *Bach2, Klf2, IL7r* and *Sell* with a concomitant increase in gene expression highlighting a role of exogenous lactate induced H3K9la in driving gene expression in naïve CD8 T cells (Figure S8H).

Overall, these data illustrate the role of H3K18la and H3K9la in CD8 T cell subsets linked to their metabolic status. In activated CD8 T cells characterized by high levels of endogenous glycolysis-derived intracellular lactate, exogenous lactate does not significantly impact histone lactylation patterns. However, in naïve/memory subsets with low glycolysis, exogenous lactate does affect histone lactylation levels. Further, unlike in activated CD8 T cells, glycolysis does not play a significant role in driving histone lactylation in naïve/memory CD8 T cells. Instead, mitochondrial metabolism pathways exert a selective impact on H3K9la in memory CD8 T cells.

### Modulation of H3K18la and H3K9la regulates CD8 T cell mediated-anti tumor immunity

Next, to delineate the functional impact of histone lactylation in CD8 T cells, we modulated the H3K18la and H3K9la levels in activated CD8 T cells by targeting the metabolic and epigenetic pathways. First, we targeted the glycolytic pathway through LDHA inhibition which led to attenuation of H3K18la and H3K9la in activated CD8 T cells (Figure 4B, Figure S7C, D). Importantly, we noted depletion of H3K18la from promoter regions of critical T cell activation genes including *Gzmb, Pdcd1* and *Ifngr1* with a concomitant reduction in the expression of these genes (Figure 5a, b, Figure S9a-c). Additionally, LDHA inhibition significantly reduced the production of effector molecules driving CD8 T cell-mediated cytotoxicity, including Granzyme B (Figure 5B, Figure S9D) and interferon-gamma (Ifng) (Figure S9E). Next, to assess the effect of LDHA inhibition on the cytotoxicity function of CD8 T cells, we co-cultured SIINFEKL primed OT-I CD8 T cells with ova expressing MB49 cancer cells (MB49-ova) in the presence or absence of LDHA inhibitor. We observed a higher ratio of cancer cells to CD8 T cells and a lower percentage of annexin-positive apoptotic cancer cells demonstrating reduction in T cell-mediated killing of cognate antigen expressing cancer cells following LDHA inhibition (Figure 5C, Figure S9F). Together, these findings demonstrate that LDHA inhibition-mediated depletion of H3K18la impairs CD8 T cell effector function.

**Figure 5:**
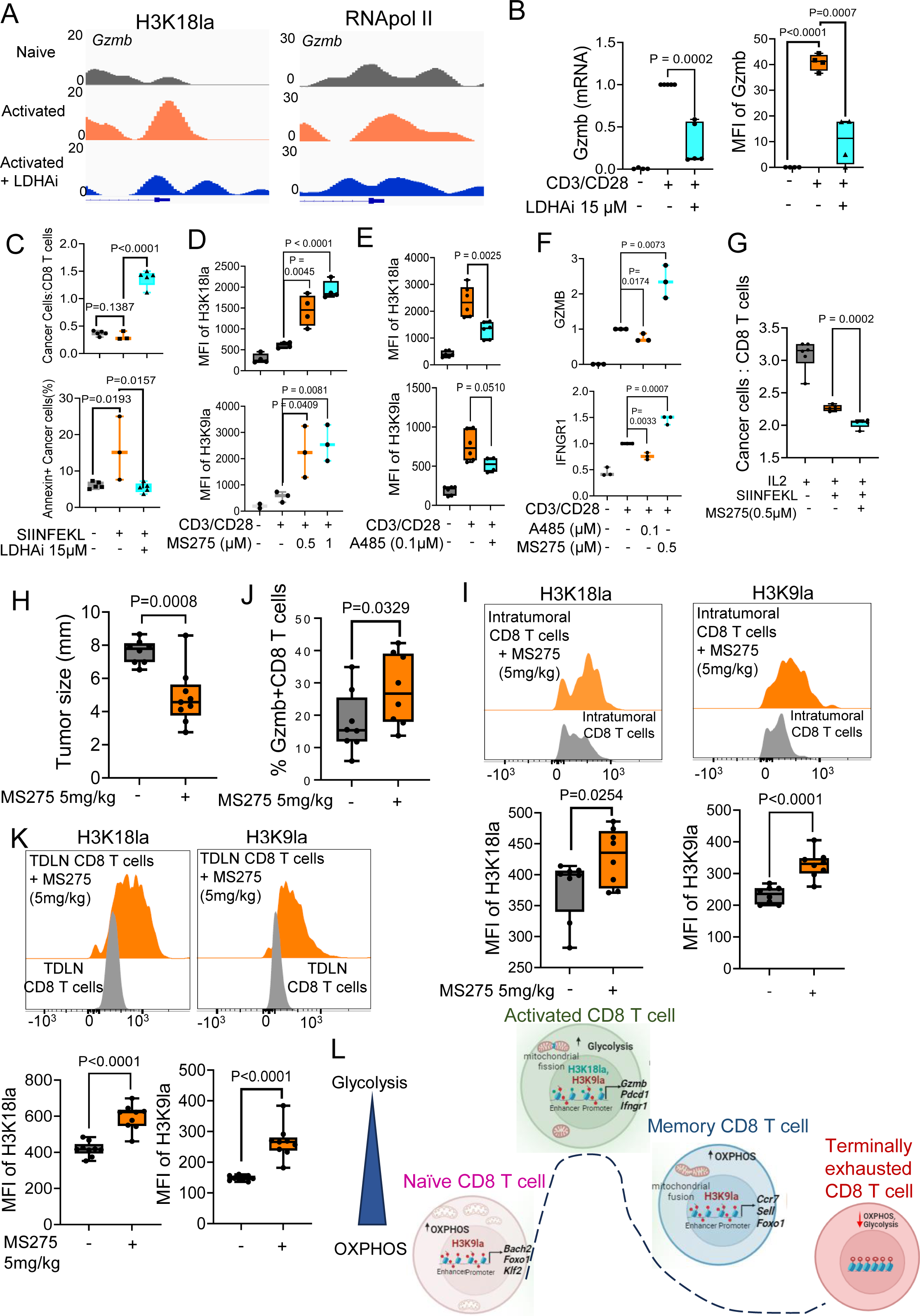
Modulation of H3K18la and H3K9la regulates CD8 T cell mediated-anti tumor immunity. **(A)** Genome Browser view of H3K18la and RNApol II peaks at the Gzmb gene promoter region (TSS ± 1kb) in the indicated CD8 T cell subsets. **(B)** Box-and-whisker plots showing the relative expression of Granzyme B (Gzmb) (Left panel) and the MFI of Gzmb (Right panel) in naïve, activated, and 15 µM LDHA inhibitor treated activated CD8 T cells. (n=4) Two-tailed Student’s t-test was performed. **(C)** Box and whisker plot representing the ratio of MB49-ova cancer cells: T cells following 24-hour co-culture of the cancer cells with unstimulated, SIINFEKL pre-stimulated and 15 µM LDHA inhibitor treated OT-I CD8 T cells (top panel). Box and whisker plot representing the percentage of annexin-V positive MB49-ova cancer cells following 24-hour co-culture with unstimulated, SIINFEKL pre-stimulated and 15 µM LDHA inhibitor treated OT-I CD8 T cells (bottom panel). n=3-5. Two-tailed Student’s t-test was performed. **(D)** Box-and-whisker plots demonstrating the MFI of H3K18la and H3K9la in naïve, activated, 0.5 µM and 1 µM MS275 (HDAC 1-3 inhibitor) treated activated CD8 T cells. (n=3) Two-tailed Student’s t-test was performed. **(E)** Box-and-whisker plots showing the MFI of H3K18la and H3K9la in naïve, activated, and 0.1 µM A485 (Ep300/CBP inhibitor) treated activated CD8 T cells. (n=6) Two-tailed Student’s t-test was performed. **(F)** Box-and-whisker plots depicting the relative expression of Granzyme B (Gzmb) and Interferon gamma receptor (IFNGR1) in naïve, activated, 0.1 µM A485 and 0.5 µM MS275 treated activated CD8 T cells. (n=3) Two-tailed Student’s t-test was performed. **(G)** Box and whisker plot representing the ratio of MB49-ova cancer cells: T cells following 24-hour co-culture of the cancer cells with unstimulated, SIINFEKL pre-stimulated and 0.5 µM MS275 treated OT-I CD8 T cells. (n=6) Two-tailed Student’s t-test was performed. **(H)** Box and whisker plot demonstrating the tumor size in untreated and 5mg/kg MS275 treated MB49 tumor-bearing mice. (n=9) One-tailed Student’s t-test was performed. **(I)** Box and whisker plot showing the relative abundance (percentage) of Granzyme B positive intratumoral CD8 T cells in untreated and 5mg/kg MS275 treated MB49 tumor-bearing mice. (n=9) One-tailed Student’s t-test was performed. **(J)** Representative histograms and Box-and-whisker plots depicting MFI of H3K18la and H3K9la in tumor derived CD8 T cells from untreated and 5mg/kg MS275 treated MB49 tumor-bearing mice. (n=9) One-tailed Student’s t-test was performed. **(K)** Representative histograms and Box-and-whisker plots demonstrating MFI of H3K18la and H3K9la in tumor draining lymph node derived CD8 T cells from untreated and 5mg/kg MS275 treated MB49 tumor-bearing mice. (n=9) One-tailed Student’s t-test was performed. **(L)** Graphical summary of the findings presented in this study depicting the role of histone lactylation in the regulation of gene transcription, phenotype and function in different CD8 T cell subsets linked to their distinct metabolic profiles. Whiskers for all box-and-whisker plots in Figure 5 represent minimum-maximum values. (See also Figure S9)

In addition to targeting metabolic enzymes to modulate histone lactylation, we directly manipulated the writer and eraser proteins responsible for regulating histone lactylation. CBP/EP300 is the known histone lactylation writer (histone lactylase) while histone deacetylases 1-3 are the known erasers for histone lactylation (histone delactylases)(Moreno-Yruela et al., 2022; Zhang et al., 2019). We treated activated CD8 T cells with the histone delactylase inhibitor MS275. Flow cytometry-based assessment of H3K18la and H3K9la levels revealed an increase in both H3K18la and H3K9la levels following inhibition of the histone delactylases (Figure 5D, Figure S9G). Conversely, treatment with the histone lactylase inhibitor A485 significantly attenuated H3K18la and H3K9la levels (Figure 5E, Figure S9H). Next, we investigated the impact of inhibition of histone lactylase (writer) and histone delactylase (eraser)-on H3K9la and H3K18la -mediated CD8 T cell effector gene expression. We observed that inhibition of histone delactylase with MS275, which increased H3K18la and H3K9la (Figure 5D), also showed concomitant increase in the expression of critical T cell effector genes including granzyme B and interferon gamma receptor (Figure 5F). Conversely, treatment with A485, a histone lactylase inhibitor, which reduced H3K18la and H3K9la (Figure 5E), also attenuated the expression of granzyme B and interferon gamma receptor (Figure 5F). We also performed *in vitro* cytotoxicity assays using MS275 treated CD8 T cells and observed a lower ratio of cancer cells: cognate T cells indicating increased tumor cell killing by MS275 treated CD8 T cells (Figure 5G). Cumulatively, these data demonstrate that inhibition of histone lactylation writer protein EP300 attenuates histone lactylation with a concurrent decrease in the expression of effector genes in activated CD8 T cells. Conversely, inhibition of histone lactylation erasers (HDACs 1-3) increases H3K18la and H3K9la with a concomitant increase in effector gene expression and CD8 T cell-mediated cytotoxicity.

Importantly, to elucidate the link between histone lactylation and T cell function *in vivo*, we treated MB49 tumor-bearing mice with MS275 (inhibitor of histone delactylase). We noted a significant reduction in tumor sizes with concurrent increase in the abundance of Gzmb+ effector CD8 T cells within the tumor in MS275 treated mice (Figure 5H, I). Notably, we also observed a significant increase in both H3K18la and H3K9la in tumor-derived and tumor draining lymph node (TDLN)-derived CD8 T cells following treatment with MS275 (Figure 5J, K). Thus, these findings underscore the impact of histone lactylation in the regulation of CD8 T cell mediated anti-tumor immunity.

Overall, this study demonstrated that lactate derived histone lactylation (H3K18la and H3K9la) marks play a pivotal role in governing the transcriptomic, metabolic, and functional profiles of CD8 T cell subsets (Figure 5L). Thus, these findings demonstrate a dynamic interplay between epigenetic and metabolic pathways in regulating CD8 T cell function including anti-tumor immunity.

## Discussion

This study uncovered the fundamental role of H3K18la and H3K9la in the regulation of CD8 T cell phenotype and function linked to their metabolic status. We noted significant enrichment of H3K18la and H3K9la in human and murine CD8 T cells, both *in vitro* and *ex vivo*, underscoring the importance of histone lactylation as an essential epigenetic modification in CD8 T cells. Further, we demonstrated the relevance of CD8 T cell histone lactylation *in vivo* utilizing two distinct disease models-GvHD and tumor. We observed that H3K18la and H3K9la play a dual role, not only marking active promoters but also actively participating in the regulation of enhancer regions. Further, we provided insight into the role of H3K18la and H3K9la as key transcription initiators in CD8 T cells either individually or in conjunction with other well established active hPTMs. These findings underscored the complex interplay of histone lactylation with other epigenetic modifications in shaping the transcriptional landscape of CD8 T cells. Importantly, we established the significance of H3K18la and H3K9la as the primary histone PTMs regulating several critical genes that control CD8 T cell phenotype and function.

Inhibition of individual cellular metabolic pathways allowed us to identify endogenous metabolic pathways, particularly glycolysis, as the primary driver of H3K18la and H3K9la in activated CD8 T cells. Unlike H3K18la, H3K9la levels remain high in both naïve and memory CD8 T cells, despite their low glycolytic activity. Our data demonstrated that mitochondrial metabolic pathways, including the TCA cycle, FAO, and ACLY selectively contribute to H3K9la in memory CD8 T cells. Of note, ETC inhibition in activated CD8 T cells increased histone lactylation possibly due to the shunting of glycolysis-derived pyruvate towards lactate generation instead of fueling the TCA cycle. Conversely, in memory CD8 T cells reliant on OXPHOS and FAO for energy generation, inhibition of ETC and FAO attenuated H3K9la, likely by disrupting the generation of key TCA cycle intermediates including oxaloacetate and malate which can fuel pyruvate and lactate production. This underscores the intricate interplay between glycolysis and mitochondrial metabolism in shaping histone lactylation patterns. Together, these data provided important insights into the distinct metabolic pathways regulating H3K18la and H3K9la marks in different CD8 T cell subsets. However, how lactate generated from distinct metabolic pathways regulates different histone lactylation marks will require further investigation.

While specific endogenous metabolic pathways drive H3K18la and H3K9la linked to the phenotypic status of CD8 T cell subsets, H3K18la and H3K9la in turn regulate key genes determining the CD8 T cell state with H3K9la playing a major role in naïve, activated, and memory CD8 T cells and H3K18la playing a critical role in activated CD8 T cells. Importantly, we found that H3K9la is closely associated with mitochondrial fusion, a process crucial for the formation of larger, interconnected mitochondria observed in naive and memory T cells(Buck et al., 2016; Lisci and Griffiths, 2023). On the other hand, H3K18la enrichment was noted on critical genes regulating mitochondrial fission, promoting the fragmentation of mitochondria, which is particularly significant in activated CD8 T cells with high glycolytic activity(Buck et al., 2016; Lisci and Griffiths, 2023). Additionally, LDHA inhibitor induced H3K18la depletion and skewing of mitochondrial morphology towards the network/diffused pattern characteristic of naïve and memory CD8 T cells indicate that targeting H3K18la can modulate mitochondrial dynamics in T cells and impact their metabolic profiles and functional states. The distinct involvement of H3K18la and H3K9la in shaping mitochondrial dynamics highlights their role as potential epigenetic regulators of CD8 T cell metabolism. By impacting mitochondrial fission and fusion, H3K18la and H3K9la likely influence the balance between glycolysis and OXPHOS, two fundamental metabolic pathways which in turn perpetuate a feed-forward loop by regulating H3K18la and H3K9la and CD8 T cell functionality.

Notably, our data demonstrate that exogenous lactate does not regulate H3K18la and H3K9la in activated CD8 T cells. However, exogenous lactate does impact H3K18la and H3K9la in naïve and memory CD8 T cells. Of note, we have used sodium lactate, a pH-neutral form of lactate, instead of lactic acid to eliminate the non-specific impact of acidosis. The differential impact of exogenous lactate on different CD8 T cell subsets may be attributed to their distinct endogenous metabolic profiles. During T cell activation, high levels of endogenous lactate production may result in the saturation of H3K18la and H3K9la levels in activated CD8 T cells, preventing further increase upon exogenous lactate addition. Additionally, MCTs which transport lactate across the cell membrane along their concentration gradient, do not import exogenous lactate into activated CD8 T cells. Instead, in activated CD8 T cells, characterized by high levels of endogenous glycolysis-derived lactate, MCTs export endogenous lactate, thus decreasing histone lactylation. Conversely, in naïve CD8 T cells with lower intracellular glycolysis-derived lactate, MCTs import exogenous lactate into the cells thereby enhancing histone lactylation. Additionally, exhausted CD8 T cells which have low endogenous glycolysis and mitochondrial metabolism(Franco et al., 2020), showed attenuation of H3K18la and H3K9la. Importantly, we noted that in terminally exhausted CD8 T cells, exhaustion-associated genes are marked by H3K9ac and not by H3K18la or H3K9la.

A critical finding of our study was the attenuation of CD8 T cell effector function following LDHA inhibitor-mediated downregulation of H3K18la and H3K9la, establishing the significance of these lactylation marks as critical regulators of T cell function. Of note, targeting metabolic enzymes including LDHA inhibition may have non-specific effects on gene expression in CD8 T cells. Therefore, we directly modulated the “writer” and “erasers” of histone lactylation to specifically demonstrate the role of histone lactylation as a pivotal regulator of gene expression and function in activated CD8 T cells.

Overall, this study demonstrates the fundamental role of histone lactylation in regulating the transcriptomic profile and function of CD8 T cells and provides insight into the distinct role of H3K18la and H3K9la in modulating CD8 T cell phenotype intricately linked to their metabolic state (Figure 5L). Recent work has also demonstrated that H4K12la enrichment of glycolytic genes regulates gene expression in microglia in a preclinical model of Alzheimer’s disease(Pan et al., 2022). It will be necessary to further explore the potential roles of H4K12la and other histone lactylation modifications in regulation of CD8 T cell homeostasis and functions. Modulating the activity of enzymes responsible for histone lactylation, or targeting specific lactylation marks, may offer novel therapeutic avenues for fine-tuning T cell responses in various disease contexts. HDAC inhibitors have received FDA approval for the treatment of hematological malignancies and are currently being tested in clinical trials for solid tumors. Our preclinical studies demonstrated that HDAC inhibition attenuated tumor size and increased effector CD8 T cell infiltration in tumors with concurrent increase in histone lactylation. Hence, the role of HDAC inhibitors in modifying histone lactylation and driving CD8 T cell mediated anti-tumor immunity in patient samples warrant further investigation. Additionally, while multiple OXPHOS inhibitors are under clinical development as cancer therapeutics, it is important to understand the nuances of globally targeting these pathways considering the differential effects of these inhibitors on cancer cells and immune cells.

## Methods

### Ethics Statement

The research outlined in this study complies with all relevant ethical regulations. Animal experiments were approved by the Animal Resource Center at The University of Texas MD Anderson Cancer Center.

### Human PBMC isolation and stimulation

Buffy coats derived from healthy human donors were collected from the blood bank. The blood was diluted 1:2 in PBS before 40ml of the diluted blood was gently layered on top of 10ml Ficoll Histopaque. The tubes were centrifuged at 1000Xg for 20 minutes at 25^ο^C in a swing bucket rotor. The white buffy coat layer containing the PBMCs present at the interphase was washed twice via centrifugation in 10ml of PBS. The pellet obtained was subjected to RBC lysis using ACK Lysis Buffer before being counted. 1.5-2X10^6^ PBMCs/well were cultured in RPMI media supplemented with 10% FBS in 96 well flat-bottomed plates at 37^ο^C and 5% CO_2_. For stimulation of CD8 T cells, the wells were coated with 2 µg/mL of anti-CD3 and 3 µg/mL of anti-CD28 was added to the culture.

### Flow cytometry

2X10^6^ cells were washed once in PBS containing 5% FBS (FACS buffer) by centrifugation at 200Xg for 5 minutes at 4 °C. Following the wash cells were incubated in blocking buffer containing bovine, murine, rat, hamster, rabbit serum and 25 μg/mL 2.4G2 antibody (Fc block) in PBS for 15 minutes on ice. Cells were then incubated for 30 minutes on ice in an antibody cocktail for surface staining (Supplementary Table 1). Following surface staining, cells were washed twice in FACS buffer before being subjected to fixation and permeabilization for 45 minutes on ice. After a wash with 1X permeabilization buffer, intracellular staining was performed by incubating the cells for 20 minutes in a cocktail of antibodies (Supplementary Table 1) at room temperature. Then the cells were washed twice in FACS buffer, fixed in 1% paraformaldehyde and acquired on a BD LSRFortessa flow cytometer.

### Mice

5–7-weeks old C57BL/6 mice, Bhlhe40 KO, Pmel-I and OT-I mice were used in this study. Age and sex matched mice were used for experiments. 10-12 weeks old female BALB/c and C57BL/6 mice were used for the GvHD experiments. Mice were housed at the Animal Resource Center, University of Texas MD Anderson Cancer Center where they were maintained in specific pathogen-free conditions, 20-25°C, 30-70% humidity and on a 12hour light/12hour dark cycle. Animal protocols were approved by the Institutional Animal Care and Use Committee of the University of Texas MD Anderson Cancer Center.

### Murine splenocyte isolation and stimulation

C57BL/6 mice, Bhlhe40 KO, Pmel-I and OT-I mice were dissected, and spleens harvested. Spleens were washed in ice-cold PBS before being mushed using the plunger of a 1ml syringe and passed through a 70µ strainer. The single cell suspension obtained was washed in PBS by centrifugation at 220Xg for 5 minutes at 4^ο^C and the cell pellet was subjected to RBC lysis using ACK Lysis Buffer before being counted. 1.5-2 X10^6^ splenocytes/well were cultured in RPMI media supplemented with 10% FBS in 96 well flat-bottomed plates at 37^ο^C and 5% CO_2_. For stimulation of CD8 T cells, the wells were coated with 3 µg/mL of anti-CD3 and 2 µg/mL of CD28 was added to the culture. For the generation of memory CD8 T cells, we stimulated activated CD8 T cells with 20ng/ml of murine recombinant IL-15 for additional 6-8 days with subculturing every 3 days. For the generation of terminally exhausted CD8 T cells, we stimulated activated CD8 T cells with 3 µg/mL of anti-CD3 plus 10ng/ml recombinant murine IL-2 in the absence of CD28 co-stimulation for 21 days with subculturing every 48 hours. For specific experiments, splenocytes were stimulated with 1 µg/mL hGP100 peptide, 1 µg/mL LCMV peptide or 9.63 µg/mL SIINFEKL peptide as outlined in the Figurelegends. Further, to test the effect of pharmacological inhibition of specific metabolic pathways on H3K18la and H3K9la, naïve, activated and memory CD8 T cells were treated with inhibitors as indicated in the Figurelegends. To test the impact of exogenous lactate on the different CD8 T cell subsets, the cells were cultured in the presence of 25mM sodium-L lactate for 24 hours.

### MB49-ova OT-1 tumor model

OT-I mice were subcutaneously injected in the right flank with 200X10^3^ MB49 control and MB49-ova cells resuspended in 100µl of PBS per mouse. On Day 10 following injection, mice were sacrificed and the tumor draining lymph nodes were isolated, mushed and a single-cell suspension was prepared as described above, for flow cytometry staining and acquisition.

### MB49 tumor model for CD8 T cell exhaustion

C57BL/6 mice were subcutaneously injected in the right flank with 200X10^3^ MB49 cells resuspended in 100µl of PBS per mouse. On Day 10 following injection, mice were sacrificed and the tumors were isolated. Following isolation, tumors were transferred to Eppendorf tubes containing 300µl of FBS free media containing DNase I (20mg/ml) and Liberase (0.66mg/ml) and chopped with scissors. The sample Eppendorfs were then placed in a thermal shaker at 1000rpm at 37°C for 30 minutes for tumor digestion. Following digestion the samples are mushed and a single-cell suspension was prepared as described above, for flow cytometry staining and acquisition.

### MB49 tumor model with *in vivo* MS275 treatment

C57BL/6 mice were subcutaneously injected in the right flank with 200X10^3^ MB49 cells resuspended in 100µl of PBS per mouse. On day 5, 7, 9 and 12 post-tumor inoculation, mice received intraperitoneal injections of 5 mg/kg of MS275 resuspended in DMSO and PBS. On Day 12 the tumors were measured with a Vernier Calliper to assess the differences in tumor sizes. Subsequently on Day 13, mice were sacrificed and the tumors and tumor draining lymph nodes were isolated. Following isolation, tumors were transferred to Eppendorf tubes containing 300µl of FBS free media containing DNase I (20mg/ml) and Liberase (0.66mg/ml) and chopped with scissors. The sample Eppendorfs were then placed in a thermal shaker at 1000rpm at 37°C for 30 minutes for tumor digestion. Following digestion, the samples were mushed and a single-cell suspension was prepared as described above, for flow cytometry staining and acquisition.

### Anti-CTLA-4 and anti-PD-1 treatment regimen

MB49 tumor-bearing C57BL/6 mice received intraperitoneal injections of 200µg, 100µg and 100µg of anti-PD1 as well as anti-CTLA-4 on day 3, 6 and 9 post tumor inoculation. On Day 14 the tumors were harvested for the preparation of a single-cell suspension and staining for flow cytometry as described above.

### GvHD model

The preclinical GvHD model was generated as reported previously(Pareek et al., 2022). Briefly, 10-12 weeks old female BALB/c mice were irradiated at a myeloablative dose of 8 Gy using 137Cs irradiator on day 0. The next day, mice intravenously received a total of 5 x 10^6^ bone marrow cells with or without 5 x 10^6^ splenocytes from age-matched donor female C57BL/6 mice. Subsequently mice were sacrificed on day 8 post-transplantation and livers were harvested for the preparation of single cell suspension and staining for flow cytometry as described above.

### Magnet Assisted Cell sorting (MACS)

MACS of CD8 T cell subsets was performed according to the manufacturer’s instructions. Briefly, murine splenocytes were resuspended in MACS buffer containing 1X PBS containing 2mM EDTA and 0.5% BSA on ice before CD8 Biotin antibody cocktail was added to each sample. The samples were resuspended well and incubated on ice for 15 minutes before anti-biotin microbeads were added, resuspended well, and incubated on ice for 20 minutes. To remove unbound antibody, samples were washed in excess MACS Buffer by centrifugation at 200Xg at 4^ο^C for 5 minutes. Next, the cell pellets were resuspended in MACS buffer, run through LS columns and washed by addition of excess MACS buffer to collect CD8 T cells. For isolation of naïve or effector CD8 T cells, the same procedure was repeated on sorted CD8 T cells using CD44+ microbeads.

### Extracellular and intracellular lactate measurement assay

To quantify the concentration of L-lactate generated following T cell activation, CD8 T cells were isolated from the spleens of C57BL/6 mice and stimulated with anti-CD3 and anti-CD28 in the presence or absence of LDHA inhibitor and POMHEX as described above. Following 6, 24, or 48 hours of stimulation, culture supernatants and cells were collected and subjected to sample processing for subsequent extracellular or intracellular lactate quantification using the L-Lactate Assay Kit according to the manufacturer’s instructions.

### ChIP Sequencing

Cells of interest were subjected to ChIP using the MAGnify Chromatin Immunoprecipitation System according to the Manufacturer’s protocol. Briefly, 37% formaldehyde was added to the samples, mixed thoroughly, and incubated at room temperature for 10 minutes. Next, 1.5 M glycine was added to the samples and incubated at room temperature for 5 minutes. Following the crosslinking step, the samples were washed in cold PBS three times by centrifugation at 200Xg for 10 minutes at 4^ο^C. Next, the cell pellets were resuspended in lysis buffer and were sonicated to shear the DNA to 150-300 kb fragments. For each immunoprecipitation reaction 10µg of H3K18lac, H3K9lac, H3K4me3, H3K9ac, H3K27ac, H3K79me2, H3K36me3, H3K27me3, and RNApol II antibodies were used and the samples were incubated overnight with the antibodies at 4^ο^C. The following day, excess antibody was washed off and the target DNA was purified and eluted. The concentration of precipitated DNA was quantified using a Qubit and dsDNA HS Assay kit. 10µg of immunoprecipitated DNA per condition was sent for sequencing to the MD Anderson Cancer Center Advanced Technology Genomics Core where sequencing was performed using the NextSeq500 instrument.

### ChIP Sequencing data analysis

The quality of CHIPseq FASTQ sequences generated as described above, were assessed using FastQC v0.11.9, followed by mapping by bowtie2 v2.3.5.1(Langmead and Salzberg, 2012) with mouse reference genome mm10. The bam files obtained from mapping were further processed using SAMBLASTER v0.1.26(Faust and Hall, 2014) and SAMTOOLS v1.13(Li et al., 2009), for duplicate removal, sorting and indexing. Further, SAMBAMBA v0.6.6(Tarasov et al., 2015) was used to normalize the bam files per read counts by performing random sampling. The genome wide annotation of aligned reads was performed using ChromHMM v1.24(Ernst and Kellis, 2017) which applies multivariate hidden Markov model (HMM) to assign states by modeling combinatorial presence and absence of each mark. The reads were subjected to “BinarizeBam” followed by “LearnModel” using mm10 assembly study enrichment and functional annotation of each mark in 7-state model. The bigwig files were generated from aligned bam reads with “bamCoverage” function of deepTools v3.5.1(Ramirez et al., 2016) and used to visualize gene tracks with Integrative Genomics Viewer (IGV) v2.16.0(Robinson et al., 2011).

The ChIP-seq signal enrichment over “Input control” background was identified using Model based analysis of ChIP-seq (MACS)2 v2.2.8(Zhang et al., 2008). The peaks with significant p-values (< 0.05) were considered for further annotation with using CHIPseeker v1.30.3(Wang et al., 2022) and clusterProfiler(Yu et al., 2012) packages in R v4.0.2. The gene ontology enrichment was performed with “enrichGO” function from clusterProfiler and list of pathways enriched in each peak set was obtained. The bed files of candidate cis-regulatory elements for pELS and dELS were obtained from the SCREEN project of ENCODE(Consortium et al., 2020) and overlapped with MACS2 peaks using “findOverlapsOfPeaks” function from ChIPpeakAnno v3.28.1(Zhu et al., 2010) package to identify enhancer regions present in respective peak-sets.

The correlation matrices for promoter, pELS and dELS peak regions between different datasets were computed with “computeMatrix” and “plotCorrelations” function of deepTools to obtain the data in “tab” formats which was plotted with “corrplot” function in R. The distribution of promoters, pELS and dELS were summarized and plotted with “plotHeatmap” function of deepTools. The pairwise combinatorial analysis was performed by finding overlapped and unique peak regions for promoters, pELS and dELS with ChIPpeakAnno package followed by Reads Per Million (RPM) estimation using EdgeR v3.36.0(Robinson et al., 2010). The command “findMotifs.pl” from Homer v4.11.1(Heinz et al., 2010) was used to identify the motifs enriched in peak regions for different marks using mm10 genome as background.

The quantitative pairwise comparisons of different datasets were performed using Manorm v1.1.4(Shao et al., 2012) using both peak (bed) and read (bam) coordinates for respective sample. The significant enrichment of target genes was estimated of the basis of M value which describes the log2 fold change and plotted with ggplot2. Differential pathways enriched among datasets were identified using GSEA v4.2.3(Subramanian et al., 2005).

### RNA sequencing

Naïve, naïve treated with exogenous lactate, activated and LDHA inhibitor treated murine activated CD8 T cells were isolated and generated as described above, washed in PBS and the cell pellets were snap-frozen in liquid nitrogen before being sent to the MD Anderson Cancer Center Biospecimen Extraction Core Facility for total RNA extraction. The RNA (a minimum of 120ng, having a RNA Integrity Number value >7), extracted from the samples was sent to the MD Anderson Cancer Center Advanced Technology Genomics for stranded paired-end mRNA sequencing on a NextSeq500 instrument.

### RNA sequencing data analysis

The paired-end raw FASTQ sequences were subjected to quality control with FastQC v0.11.9, followed by adapter removal using Trimmomatic v0.39(Bolger et al., 2014). The trimmed fastq reads were mapped to mm10 reference genome using STAR v2.7.3a(Dobin et al., 2013) and resulting bam reads were sorted and indexed with SAMTOOLS v1.13. The gene counts for each sample was estimated with HTseq v2.0.2(Putri et al., 2022) and the counts were converted to “counts per million (CPM)” for differential analysis between samples with EdgeR v3.36.0(Robinson et al., 2010). The regulatory potential scores of Chipseq peaks of different histone marks for expressed genes were calculated by integrating Chipseq experiments and RNA-seq differential expression data using BETA v1.0.7(Wang et al., 2013). The heatmaps were plotted with ggplot2 using R v4.0.2.

### Seahorse assay

Agilent Seahorse XF ATP Real-Time rate assay was done to assess the cellular metabolic activity and measure the rate of ATP production from glycolytic and mitochondrial metabolism of naïve and activated CD8 T cells according to the manufacturer’s instructions. Briefly, naïve cells and activated CD8 T cells treated as indicated in the Figurelegend, were resuspended seeded on Cell-Tak coated wells of a XF-96 well-plate, before being subjected to the XF-96 Analyzer for the measurement of the extracellular acidification rate (ECAR) and the oxygen consumption rate (OCR) under basal conditions and following sequential injections of oligomycin, FCCP and Rotenone. Real-time measurements of ATP production rates were obtained and analyzed using Seahorse Wave software.

### LDHA knockdown in primary murine CD8 T cells

The Ldha Mouse shRNA Plasmid was used to knockdown the *Ldha* gene according to the manufacturer’s instructions. Briefly, 5-10 million murine CD8 T cells sorted by MACS from the spleens of C57BL/6 mice as described above, were seeded in 2ml of complete RPMI per well in a 6 well plate. Subsequently, a mixture containing 1µg of plasmid DNA and Turbofectin transfection reagent in 250ul of Opti-MEM media was added dropwise to the CD8 T cell culture. Following 48 hours of incubation, CD8 T cells were stimulated with anti-CD3 plus anti-CD28 for 48 hours and subjected to staining and flow-cytometry acquisition as described above, to determine the impact of *Ldha* knockdown on histone lactylation levels.

### RNA Isolation and cDNA preparation for qPCR

Single cell suspension of mouse splenocytes were isolated, stimulated, treated, and sorted as described above to obtain the cell subsets of interest. The cells were resuspended in TriZol reagent before one-fifth volume of chloroform was added to the samples. Following the addition of chloroform, the samples were mixed vigorously and incubated at room temperature for 10 minutes. The samples were then centrifuged at 12,000Xg for 10 minutes at 4^ο^C. The top aqueous layer was collected, and equal volume of isopropanol was added before incubation at room temperature for 10 minutes. The samples were again centrifuged at 12000Xg for 10 minutes at 4^ο^C before a final wash by centrifugation in 75% ethanol. Finally, the pellets were air-dried and resuspended in nuclease-free water. RNA concentration was measured using a Nanodrop and 1µg of RNA was reverse transcribed into cDNA using the Superscript III cDNA kit. This cDNA was used to perform real time PCR (qPCR) to measure the expression of the genes of interest. The primer sequences used have been outlined in Supplementary Table 1.

### ChIP-qPCR

Memory CD8 T cells treated with or without 5µM of BMS-303141 were subjected to ChIP using the anti-H3K9la antibody as described above. 100ng of the immunoprecipitated DNA from each of the conditions was subjected to qPCR analysis as described above. The primer sequences used have been outlined in Supplementary Table 1.

### Enzyme-linked immunosorbent assay (ELISA)

96-well binding plates were coated overnight at 4^ο^C with mouse Ifng capture antibody, diluted 5:1000 in coating buffer containing NaHCO_3_ and Na_2_CO_3_ in deionized water. Next day, the plates are washed three times with wash buffer containing PBS + 0.05% Tween-20 before the wells were blocked with 1X assay diluent for 1 hour at room temperature with gentle agitation. Following three more washes Ifng standards and samples were added to the designated wells and incubated at room temperature for 2 hours with gentle agitation. Following incubation, the wells were washed three times and then 5:1000 diluted mouse Ifng detection antibody was added to each of the wells before incubation for 1 hour at room temperature. Following three more washes, 1:1000 Avidin-HRP solution was added to each well and incubated at room temperature for 30 minutes. For the final step, the wells were soaked in wash buffer for 30 seconds for a total of four washes before TMB substrate solution was added and incubated in the dark until color development. Reactions were then stopped by the addition of Stop Solution to each well and the optical density values were read on a microplate spectrophotometer, at 450nm. The standard curve was prepared using linear regression based on the standard absorbance values of known concentrations. This allowed for proper quantification of the concentration of Ifng present in the supernatants of the samples.

### T cell cytotoxicity assay

Single-cell suspension was prepared from spleens were harvested from OT-I mice as described above. Splenocytes were either left unstimulated or stimulated with the SIINFEKL peptide for 48 hours in the presence or absence of 15µM LDHA inhibitor or 0.5µM MS275. MB49-ova cancer cells were also seeded in 24 well plates. Following stimulation of the splenocytes for 48 hours, CD8 T cells were sorted from each treatment condition using magnet-assisted cell sorting and co-cultured with the MB49-ova cancer cells. Following 24 hours of co-culture, the cells were harvested and stained with an antibody cocktail containing CD45+ AF532, CD3e FITC, CD8a BV711, and annexin PB diluted in 1X Annexin Binding Buffer at room temperature for 15 minutes. The stained cells were then washed with 1X Annexin Binding Buffer and acquired in a BD LSRFortessa flow cytometer. FlowJo v10 was used for analysis of the flow cytometry data.

### Super-Resolution Structured Illumination Microscopy

CD44-naïve CD8 T cells were sorted from splenocytes derived from C57BL/6 mice. Additionally, splenocytes isolated from C57BL/6 mice were stimulated with anti-CD3 plus anti-CD28 in the presence or absence of 15µM LDHA inhibitor as discussed above. Following 48 hours of stimulation, CD44+ activated CD8 T cells were sorted from the stimulated splenocytes using MACS. The sorted cells were then seeded on poly Lysine coated cover slips before being stained with MitoTracker Red CMXROS and DAPI according to the manufacturer’s protocol. The stained coverslips were mounted in ProLong Gold Antifade medium before being imaged in a OMX Blaze V4 Microscope.

### Transmission electron microscopy

CD44-naïve CD8 T cells were sorted from splenocytes derived from C57BL/6 mice. Additionally, splenocytes isolated from C57BL/6 mice were stimulated with anti-CD3 plus anti-CD28 in the presence or absence of 15µM LDHA inhibitor as discussed above. Following 48 hours of stimulation, CD44+ activated CD8 T cells were sorted from the stimulated splenocytes using MACS. Sorted cells were fixed with a solution containing 3% glutaraldehyde plus 2% paraformaldehyde in 0.1 M cacodylate buffer, pH 7.3, then washed in 0.1 M sodium cacodylate buffer and treated with 0.1% Millipore-filtered cacodylate buffered tannic acid, postfixed with 1% buffered osmium tetroxide, and stained en bloc with 1% Millipore-filtered uranyl acetate. The samples were dehydrated in increasing concentrations of ethanol, infiltrated, and embedded in LX-112 medium. The samples were polymerized in a 60^ο^C oven for approximately 3 days. Ultrathin sections were cut in a Leica Ultracut microtome, placed on formvar coated single slot copper grids, stained with uranyl acetate and lead citrate and examined in a JEM 1010 transmission electron microscope at an accelerating voltage of 80 kV. Digital images were obtained using AMT Imaging System.

### Statistics & Reproducibility

R v4.0.2 and GraphPad Prism software v9 was used for the statistical analyses. The specific statistical tests used have been indicated in the respective Figure legends.

### Data and code availability

The CHIP-seq and RNA-seq data supporting this study have been deposited in Sequence Read Archive (SRA) under accession code PRJNA1040830. All requests for data should be made to the corresponding author (S.G), following verification of any intellectual property or confidentiality obligations.

## Acknowledgments

This research is supported by the MD Anderson Physician Scientist Award (S.G) and NIH-R01 Merit Award (R37 CA279192-01) (S.G). Dr. Goswami is a member of the James P. Allison Institute. We acknowledge the UT MD Anderson Center ATGC Core (grant CA016672) for the ChIP-sequencing and RNA-sequencing studies. We thank the UT MD Anderson Center Metabolomics core for the Seahorse XFe96 Flux Analyzer service, High Resolution Electron Microscopy Facility (CCSG grant NIH P30CA016672) and MDACC Advanced Microscopy Core for super-resolution microscopy imaging.

## Author Contributions

D.R and P.S designed and performed the experiments, analyzed data and wrote the manuscript. M.H performed experiments and wrote the manuscript. B.C and A.J.T helped with the murine experiments. J.S. I and A.T provided the GvHD preclinical model. S.G. developed the project, designed the experiments, analyzed data, wrote the manuscript and acquired funding.

## Supplemental information

### Supplementary Figure legends

**Figure S1:**
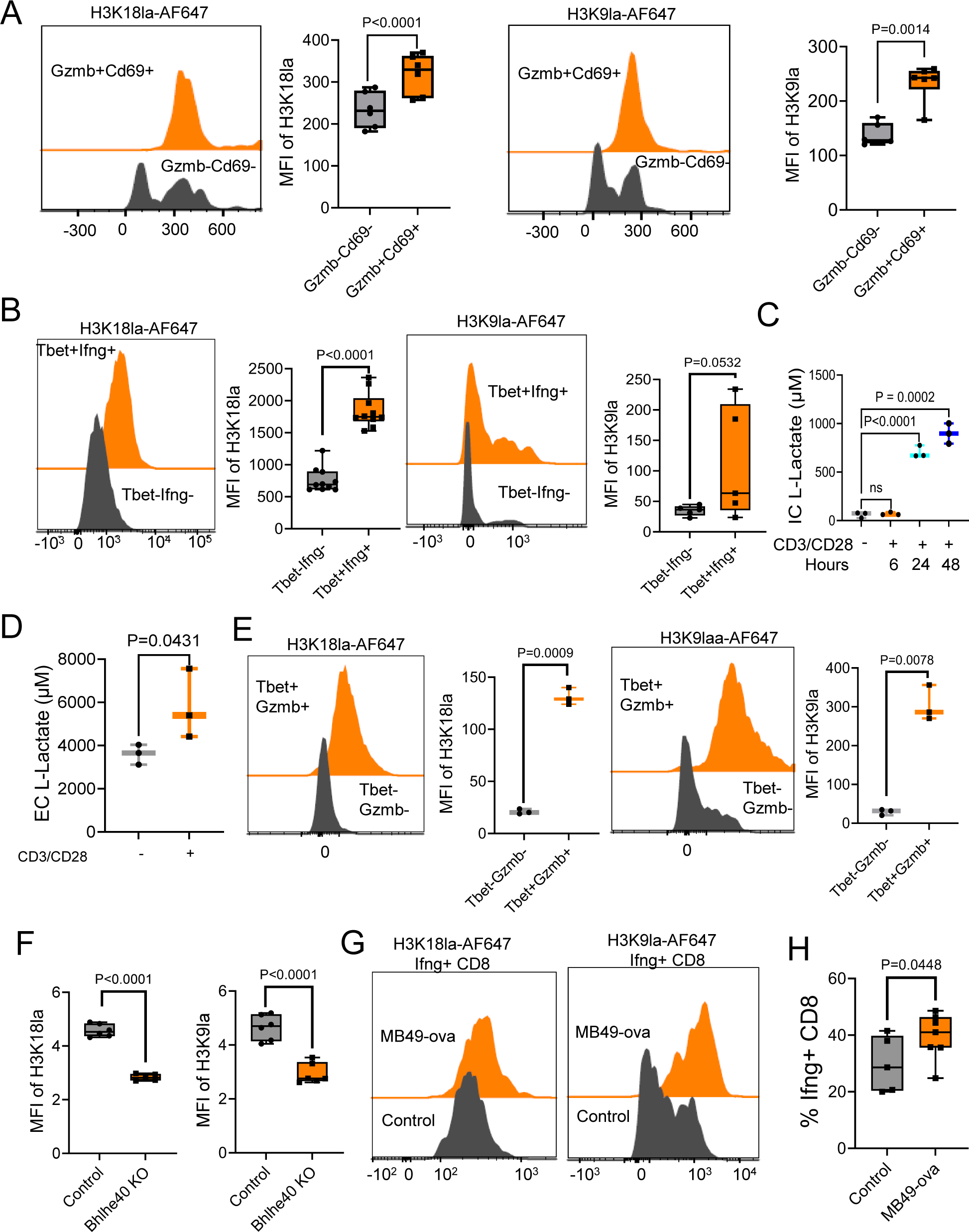
**A,** Representative histograms and box-and-whisker plots demonstrating the differences in H3K18la and H3K9la levels between human Gzmb+CD69+ and Gzmb-CD69-CD8 T cells following 24 hours of anti-CD3 and anti-CD28 stimulation. n=6 human donors. Two-tailed Student’s t-test was performed. **B**, Representative histograms and box-and-whisker plots showing H3K18la and H3K9la levels in murine Tbet+Ifng+ and Tbet-Ifng-CD8 T cells following 24 hours of anti-CD3 and anti-CD28 stimulation. n= 3 female mice. Two-tailed Student’s t-test was performed. **C**, Box and whisker plot representing the concentration of intracellular (IC) L-Lactate in naïve and activated murine CD8 T cells following 6, 24, and 48 hours of CD3/CD28 stimulation. (n=3) Two-tailed Student’s t-test was performed. **D**, Box-and-whisker plot demonstrating the concentration of L-lactate in the culture supernatant (EC) of murine CD8 T cells treated for 24 hours as indicated. n= 3 female mice. One-tailed Student’s t-test was performed. **E,** Representative histograms and box-and-whisker plots depicting H3K18la and H3K9la levels in pmel mice-derived Tbet+Gzmb+ and Tbet-Gzmb-CD8 T cells following stimulation with the cognate GP100 peptide. n= 3 female mice. Two-tailed Student’s t-test was performed. **F**, Box-and-whisker plots demonstrating H3K18la and H3K9la levels in CD8 T cells derived from control C57BL/6 mice and *Bhlhe40* KO mice following anti-CD3 and anti-CD28 stimulation for 48 hours. n=6 female mice. Two-tailed Student’s t-test was performed. **G,** Representative histograms showing H3K18la and H3K9la levels in Ifng+ CD8 T cells derived from the TDLNs of control and MB49-ova tumor-bearing mice. **H,** Box-and-whisker plots demonstrating the percentage of Ifng+CD8 T cells in TDLNs of control and MB49-ova tumor-bearing mice. n=5 female mice bearing control MB49 tumors and 7 female mice bearing MB49-ova tumors. Whiskers for all box-and-whisker plots in figure S1 represent minimum-maximum values.

**Figure S2:**
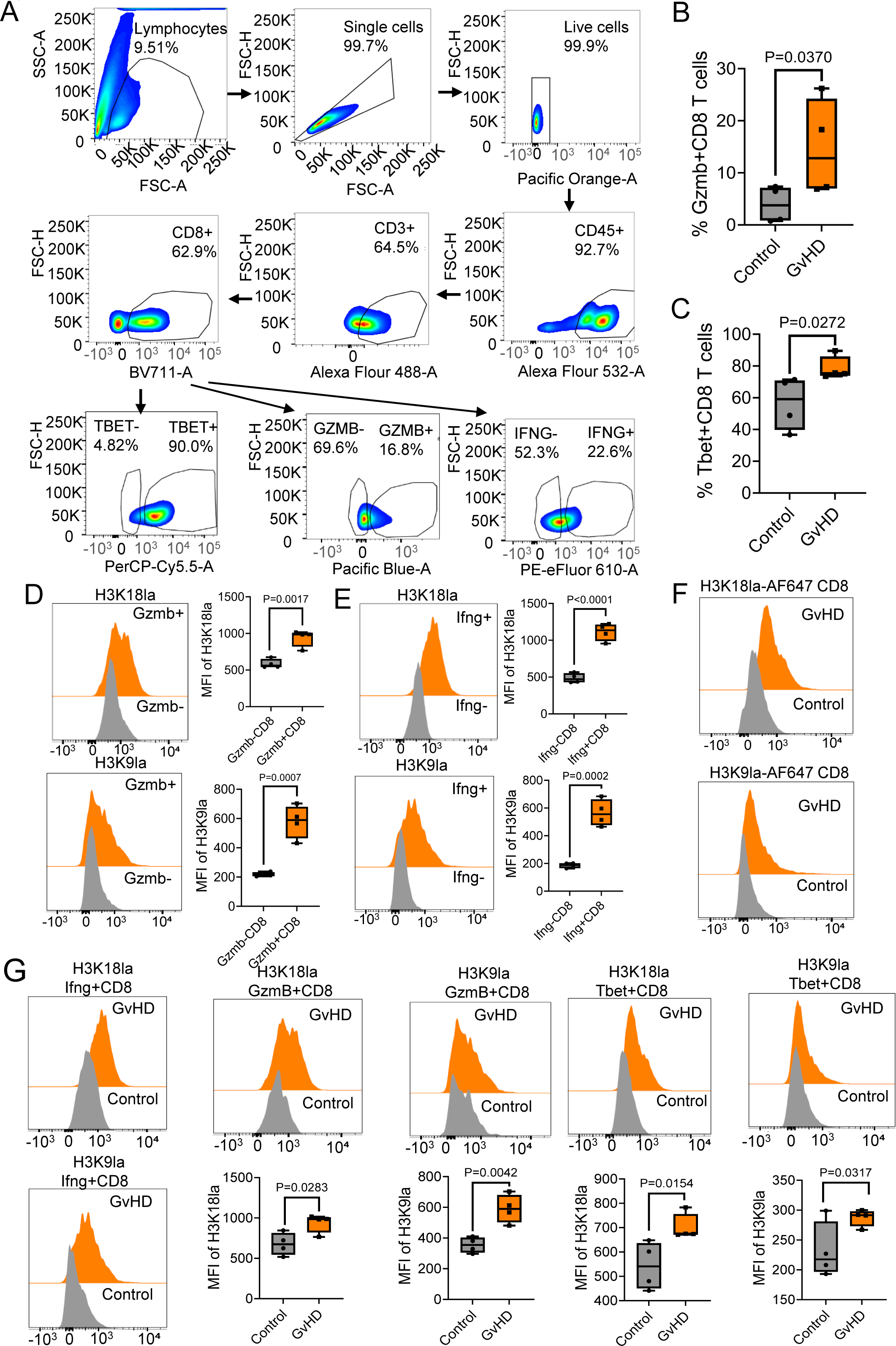
**A**, Pseudocolor plots demonstrating the gating strategy used to analyze the CD8 T cell populations in livers from control and GvHD mice. **B**, Box-and-whisker plot demonstrating the percentage of Gzmb+CD8 T cells in the livers of control and GvHD mice. n= 4 control female mice and 4 GvHD female mice. One-tailed Student’s t-test was performed. **C**, Box-and-whisker plot depicting the percentage of Tbet+ CD8 T cells in the livers of control and GvHD-afflicted mice. n= 4 control female mice and 4 GvHD female mice. One-tailed Student’s t-test was performed. **D,** Representative histograms and box-and-whisker plots demonstrating the level of H3K18la and H3K9la in Gzmb+ and Gzmb-CD8 T cells derived from the livers of GvHD mice. n= 4 GvHD-afflicted female mice. Two-tailed Student’s t-test was performed. **E,** Representative histograms and box-and-whisker plots demonstrating the level of H3K18la and H3K9la in Ifng+ and Ifng-CD8 T cells derived from the livers of GvHD mice. n= 4 GvHD-afflicted female mice. Two-tailed Student’s t-test was performed. **F**, Representative histograms showing H3K18la and H3K9la levels in murine CD8 T cells derived from the livers of control and GvHD mice. n= 4 control female mice and 4 GvHD female mice. **G**, Representative histograms and box-and-whisker plots demonstrating the level of H3K18la and H3K9la in Gzmb+, Tbet+ and Ifng+ CD8 T cells derived from the livers of GvHD (from panel d and e) and control mice. n= 4 control female mice and 4 GvHD female mice. One tailed and two-tailed Student’s t-test were performed. Whiskers for all box-and-whisker plots in figure S2 represent minimum-maximum values.

**Figure S3:**
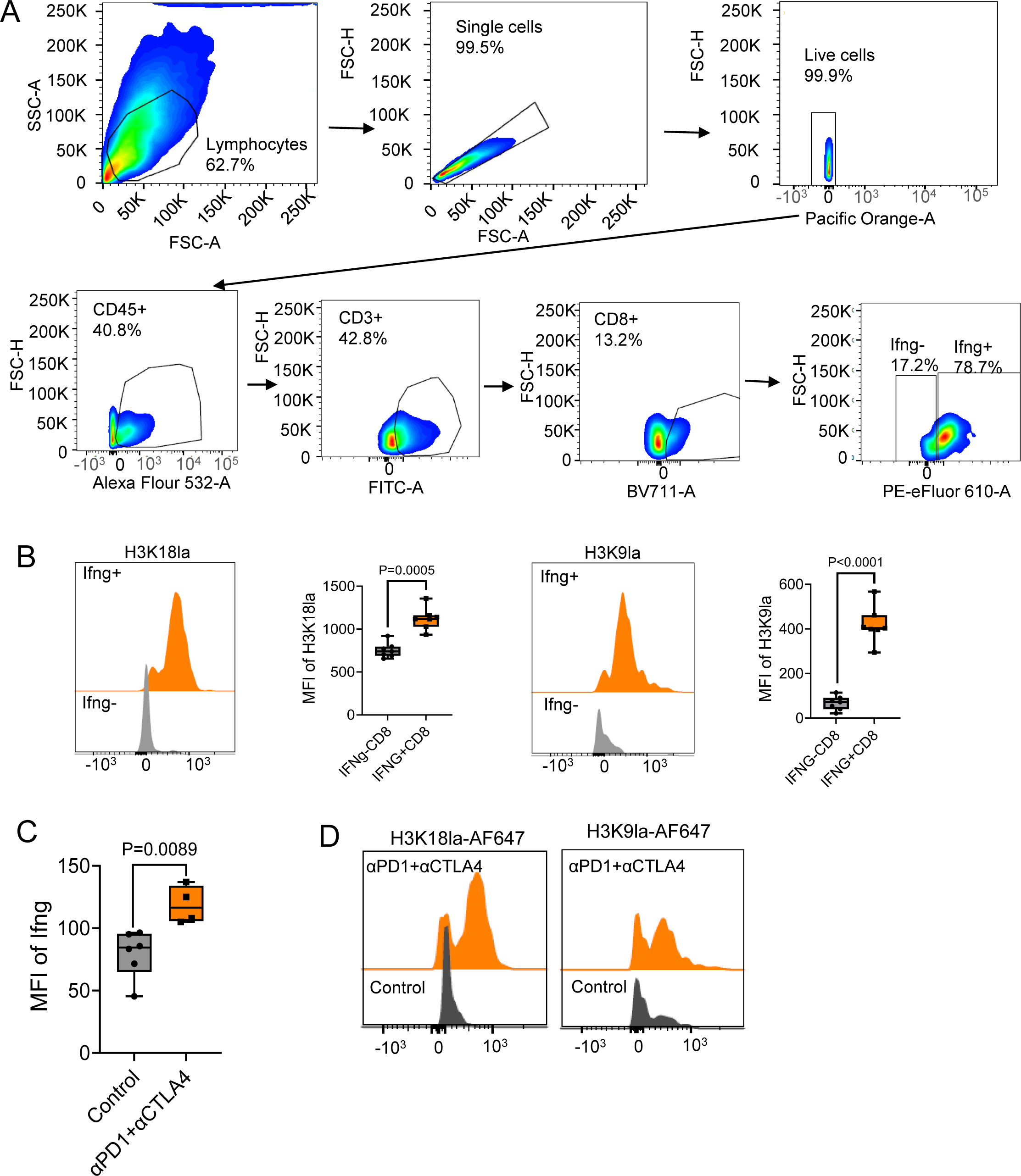
**A**, Pseudocolor plots demonstrating the gating strategy used to analyze the CD8 T cell populations in MB49 tumors. **B**, Representative histograms and box-and-whisker plots demonstrating the level of H3K18la and H3K9la in Ifng+ and Ifng-CD8 T cells derived from MB49 tumors. n= 7 female MB49 tumor-bearing mice. Two-tailed Student’s t-test was performed. **C**, Box-and-whisker plot depicting the expression of Ifng in intratumoral CD8 T cells derived from control and anti-PD1 plus anti-CTLA4 treated MB49 tumor-bearing mice. n= 6 control female mice and 4 female mice treated with a combination of anti-PD1 and anti-CTLA4. Two-tailed Student’s t-test was performed. **D**, Representative histograms demonstrating the expression of H3K18la and H3K9la in intratumoral CD8 T cells derived from control and anti-PD1 plus anti-CTLA4 treated MB49 tumor-bearing mice. Whiskers for all box-and-whisker plots in figure S3 represent minimum-maximum values.

**Figure S4:**
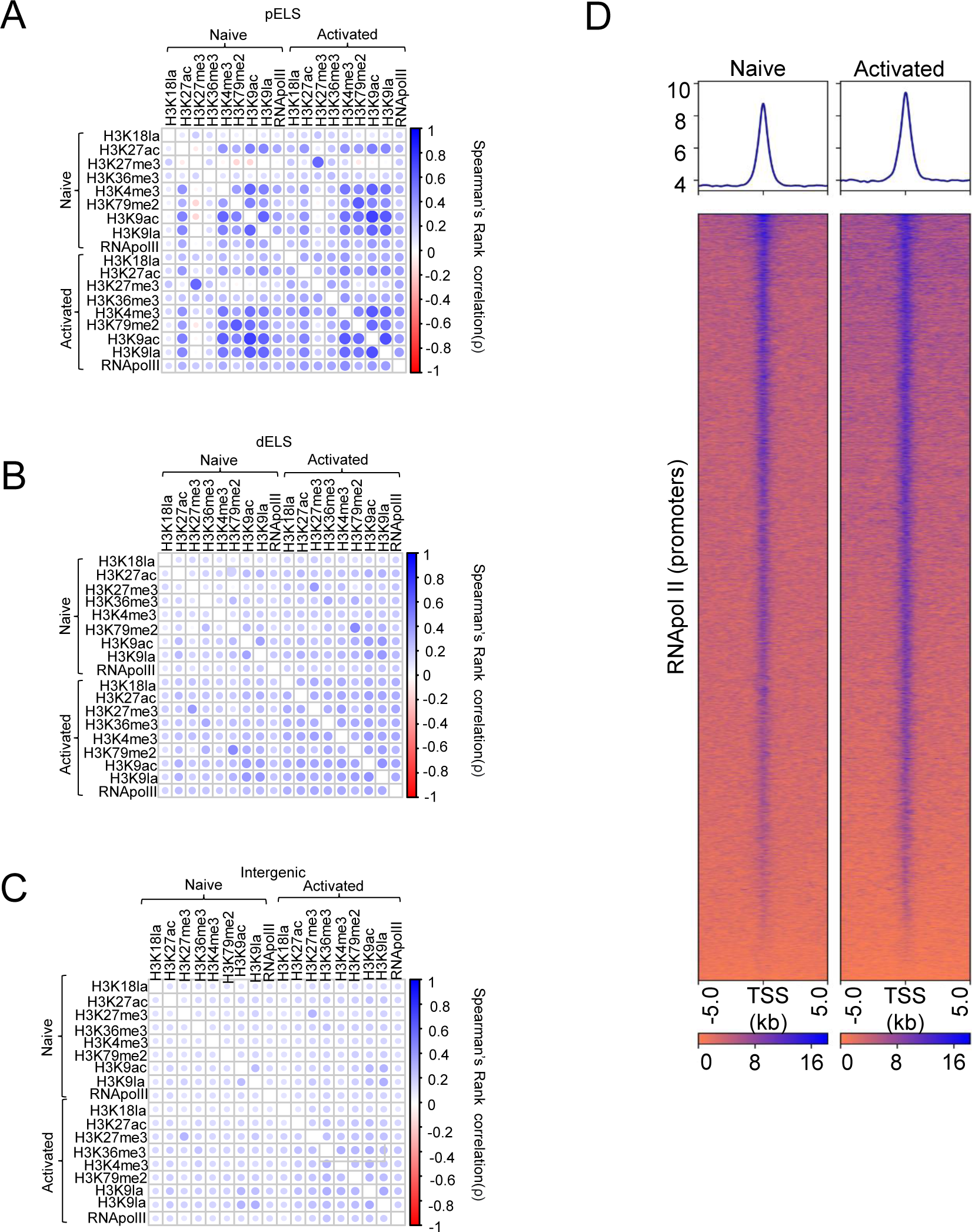
**A-C**, Correlation matrices showing the Spearman’s correlation coefficients (color intensities and the size of circles) among H3K18la and H3K9la sites and the other indicated hPTMs at proximal enhancer-like sequences (pELS) (**A**), distal enhancer-like sequences (dELS) (**B**) and intergenic regions (**C**). **D**, Heatmap depicting the genomic occupancy of RNApol II at gene promoter regions in murine naïve and activated CD8 T cells.

**Figure S5:**
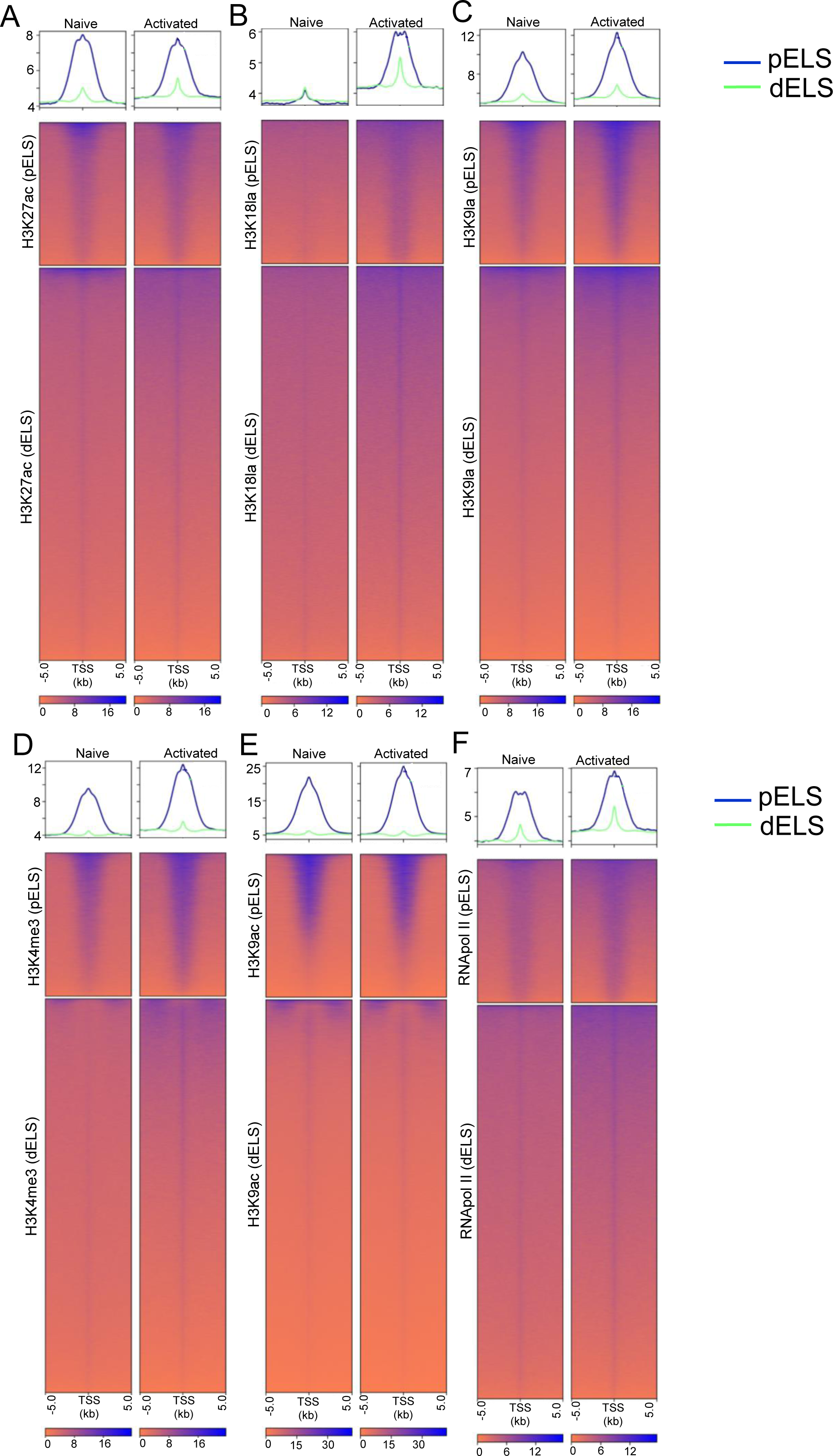
**A-F**, Heatmaps depicting the genomic occupancy of the indicated hPTMs **(A-E)** and RNApol II **(F)** at pELS and dELS in naïve and activated murine CD8 T cells.

**Figure S6:**
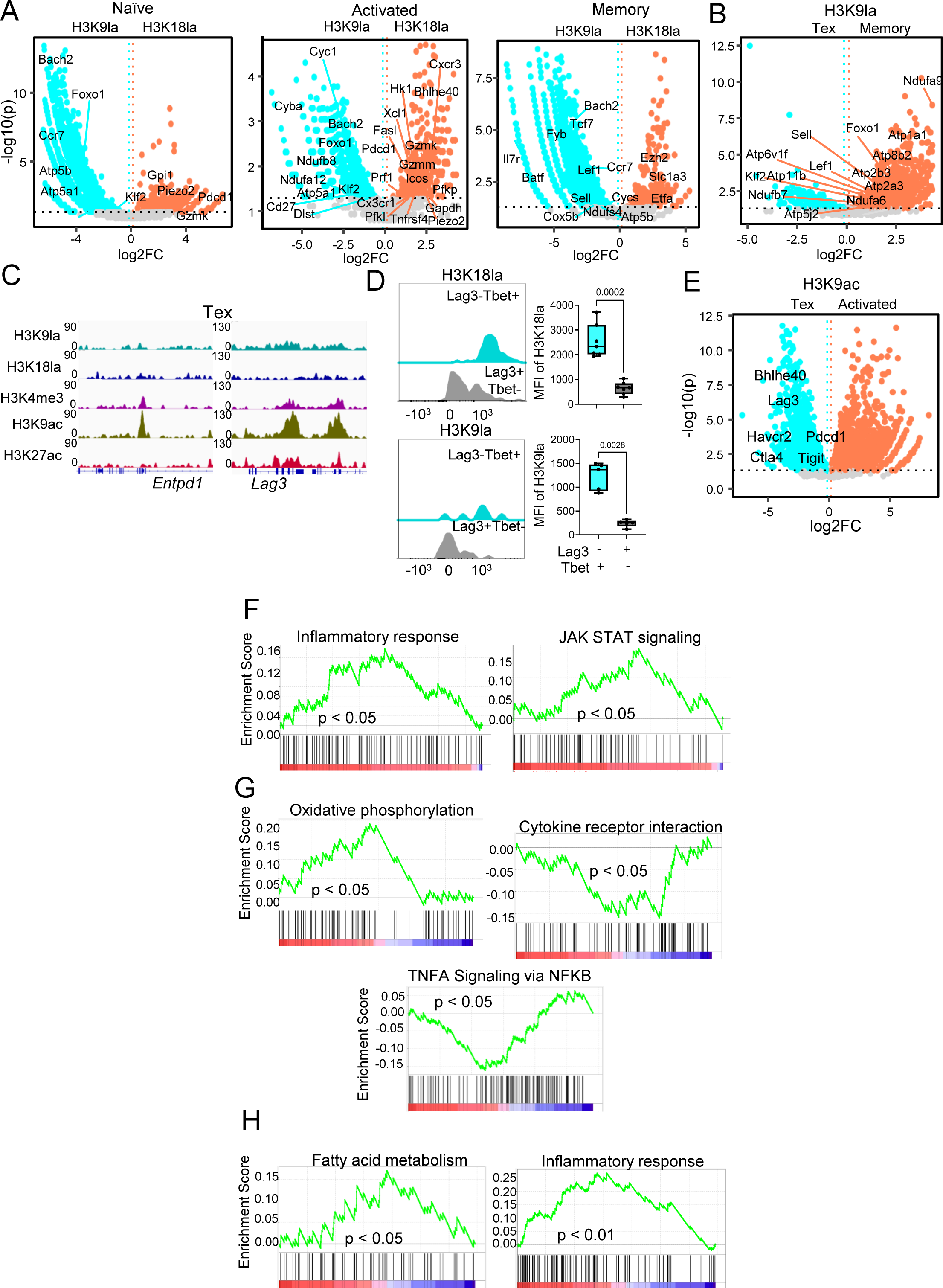
**A,** Volcano plots representing genes harboring differentially enriched H3K18la and H3K9la marks in the indicated CD8 T cell subsets. Volcano Plot shows the log2 ratio of fold change (log2FC) plotted against the Absolute Confidence, -log10 adjusted p-value (-log10(p)). The p-value was derived from the Student’s t-test using MAnorm. **B**, Volcano plot representing genes harboring differentially enriched H3K9la marks in terminally exhausted (Tex) versus memory CD8 T cells. Volcano plots show the log2 ratio of fold change (log2FC) plotted against the Absolute Confidence, -log10 adjusted p-value (-log10(p)). The p-value was derived from the Student’s t-test using MAnorm. **C**, Genome browser view of H3K9la, H3K18la, H3K4me3, H3K9ac and H3K27ac peaks for Entpd1 and Lag3 gene loci in terminally exhausted CD8 T cells generated *in vitro*. **D**, Representative histograms and box-and-whisker plots depicting MFI of H3K18la (n=7) and H3K9la (n=5) levels in intratumoral Lag3-Tbet+ and Lag3+Tbet-CD8 T cells from MB49 tumor bearing mice. Two-tailed Student’s t-test was performed. **E,** Volcano plot representing genes harboring differentially enriched H3K9ac marks in terminally exhausted (Tex) versus activated CD8 T cells. Volcano plots show the log2 ratio of fold change (log2FC) plotted against the Absolute Confidence, -log10 adjusted p-value (-log10(p)). The p-value was derived from the Student’s t-test using MAnorm. **F-H**, Plots representing GSEA pathways generated from genes with H3K18la enrichment in activated CD8 T cells compared to naïve CD8 T cells **(F)**, genes with H3K9la enrichment or depletion in activated CD8 T cells compared to naïve CD8 T cells **(G)** and genes with H3K9la enrichment in memory CD8 T cells compared to activated CD8 T cells **(H)**. P values were calculated by permutation using the GSEA software. Whiskers for all box-and-whisker plots in figure S6 represent minimum-maximum values.

**Figure S7:**
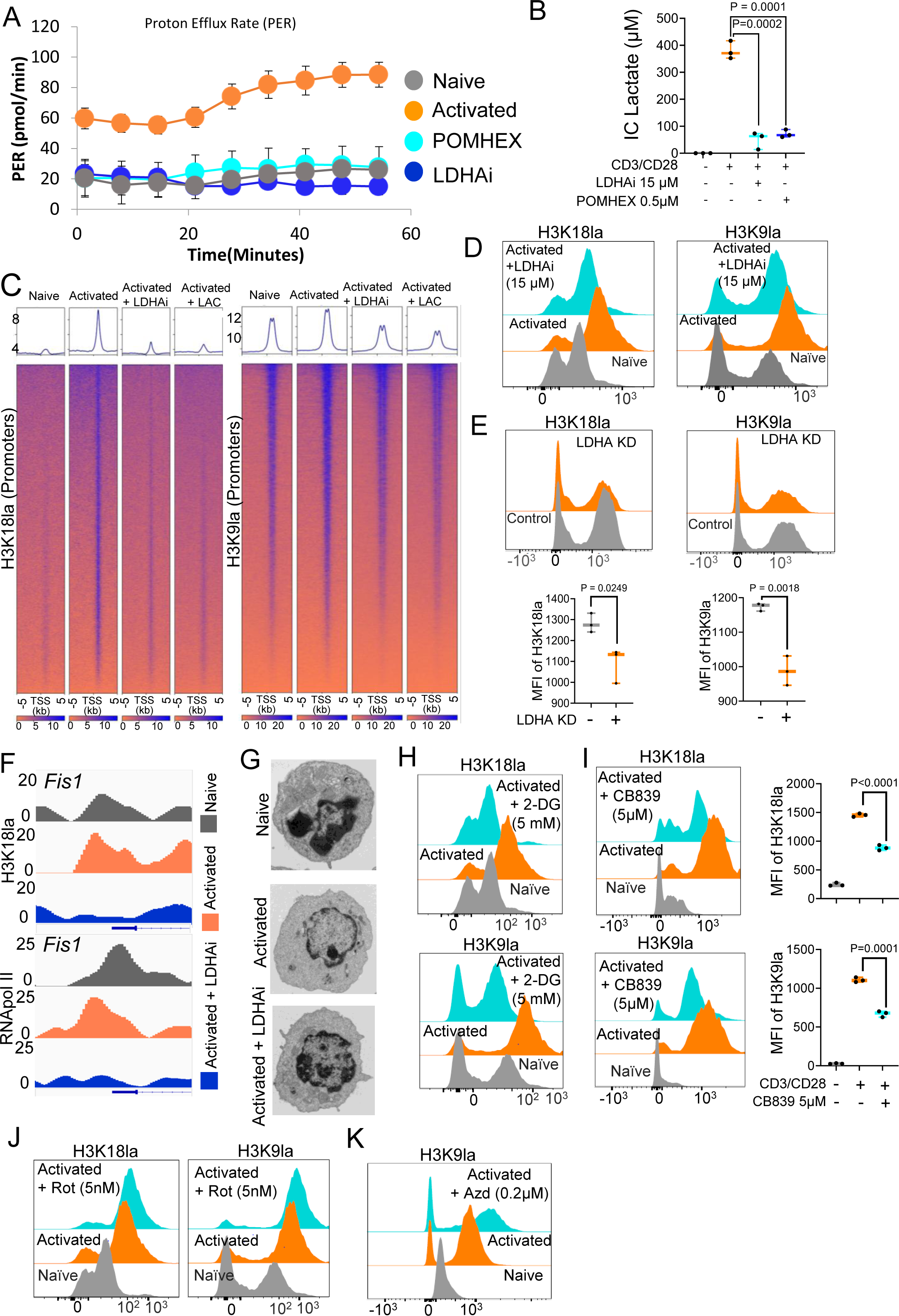
**A**, Line plot showing Proton Efflux Rate (PER) in naïve, activated, 15µM LDHA inhibitor and 0.5µM POMHEX treated activated CD8 T cells measured in an ATP Rate Determination Assay using a Seahorse XF Analyzer. **B**, Box and whisker plot representing the concentration of intracellular (IC) L-Lactate in naïve, activated, 15 µM LDHA inhibitor and 0.5 µM POMHEX treated activated CD8 T cells. (n=3) Two-tailed Student’s t-test was performed. **C**, Heatmaps indicating the genomic occupancy of H3K18la and H3K9la at TSS ± 1kb flanking TSSs in naïve, activated (from Figure 2e), 15 µM LDHA inhibitor treated (LDHA) and 25mM sodium-lactate treated (Na-La) CD8 T cells. **D**, Representative histograms depicting MFI of H3K18la and H3K9la in naïve, activated and 15 µM LDHA inhibitor treated activated CD8 T cells. **E**, Representative histograms and Box-and-whisker plots demonstrating MFI of H3K18la and H3K9la in control and *ldha* knockdown (KD) activated CD8 T cells. (n=3) Two-tailed Student’s t-test was performed. **F,** Genome Browser view of H3K18la and RNApol II binding at the *Fis1* gene promoter (TSS ±1kb) in naïve, activated (from Figure 3f) and LDHA inhibitor treated (LDHAi) activated CD8 T cells. **G,** Representative transmission electron microscopy images of naïve, activated and LDHA inhibitor(LDHAi) treated activated CD8 T cell subsets depicting mitochondrial morphology. **H,** Representative histograms showing MFI of H3K18la and H3K9la in naïve, activated and 5 mM 2-DG treated activated CD8 T cells. **I,** Representative histograms and Box-and-whisker plots indicating MFI of H3K18la and H3K9la in naïve, activated and 5 µM CB839 treated activated CD8 T cells. (n=3) Two-tailed Student’s t-test was performed. **J,** Histograms representing MFI of H3K18la and H3K9la in naïve, activated and 5 nM Rotenone treated activated CD8 T cells. **K,** Representative histogram showing MFI of H3K9la in naïve, activated and 0.2 µM Azd treated activated CD8 T cells. Whiskers for all box-and-whisker plots in figure S7 represent minimum-maximum values.

**Figure S8:**
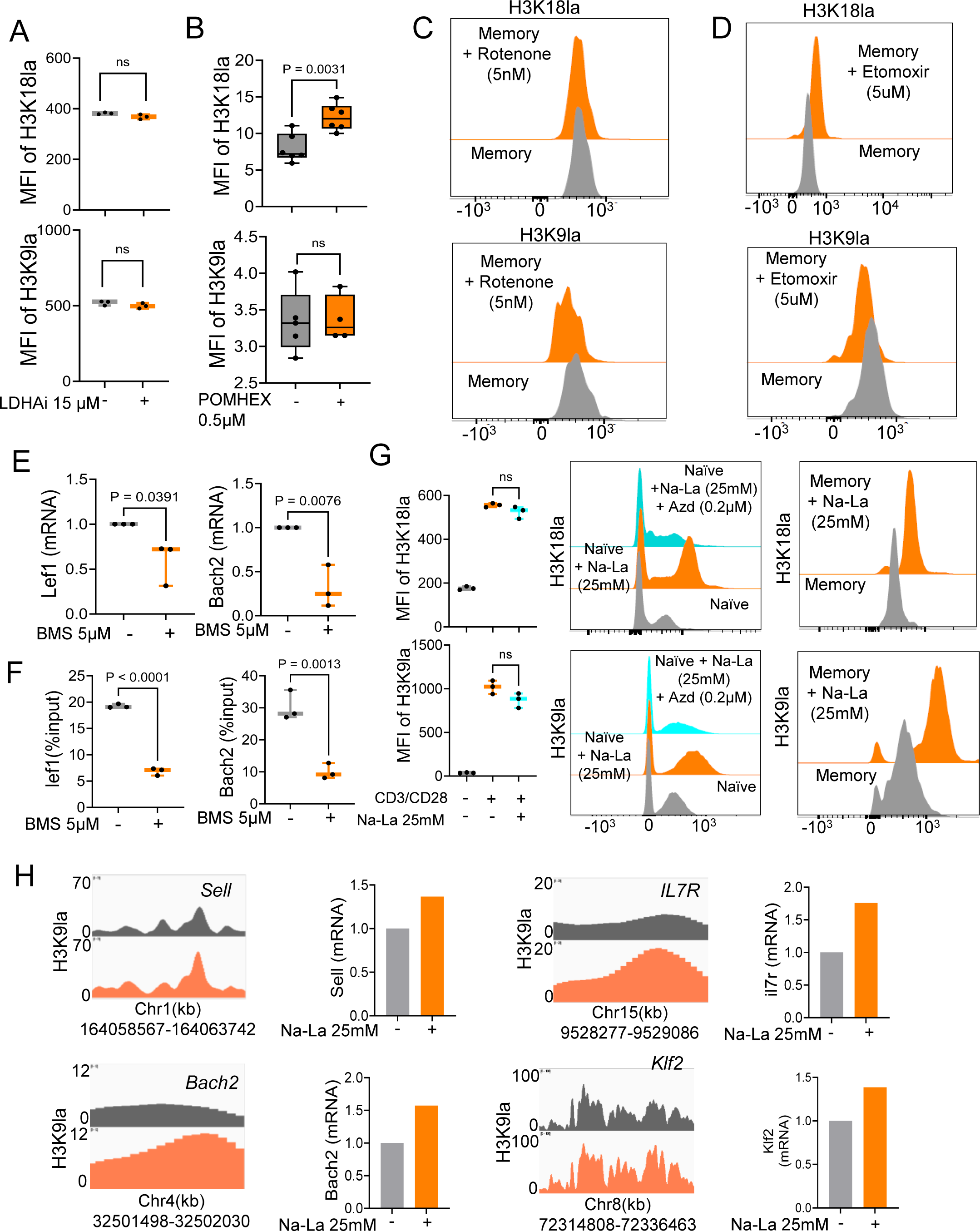
**A,** Box-and-whisker plots demonstrating MFI of H3K18la and H3K9la in untreated and 15 µM LDHA inhibitor treated memory CD8 T cells. (n=3) Two-tailed Student’s t-test was performed. ns= non-significant. **B**, Box-and-whisker plots depicting MFI of H3K18la and H3K9la in untreated and 0.5 µM POMHEX treated memory CD8 T cells. (n=6) Two-tailed Student’s t-test was performed. ns= non-significant. **C**, Representative histograms indicating MFI of H3K18la and H3K9la in untreated and 5 nM Rotenone inhibitor treated memory CD8 T cells. **D**, Representative histograms demonstrating MFI of H3K18la and H3K9la in untreated and 5 µM Etomoxir inhibitor treated memory CD8 T cells. **E**, Box-and-whisker plots indicating the relative expression of *Lef1*, and *Bach2* (mRNA) in untreated and 5 µM BMS treated memory CD8 T cells. (n=3) Two-tailed Student’s t-test was performed. **F**, Box and whisker plot showing the ChIP qPCR analysis of H3K9la enrichment at the *Lef1* and *Bach2* gene promoters in memory CD8 T cells treated with or without 5µM of the ACLY inhibitor-BMS. Data is represented as percentage of input. (n=3) Two-tailed Student’s t-test was performed. **G,** Box-and-whisker plots representing MFI of H3K18la and H3K9la in naïve, activated and 25mM sodium lactate (Na-La) treated activated CD8 T cells (left panel). (n=3) Two-tailed Student’s t-test was performed. ns= non-significant. Histograms representing MFI of H3K18la and H3K9la in naïve, 25 mM Na-La and 0.2 µM Azd (MCT1 inhibitor) treated naïve CD8 T cells (middle panel). Representative histograms showing the MFI of H3K18la and H3K9la in memory CD8 T cells treated with and without 25mM Na-La (right panel). **H,** Genome browser plots and bar plots of H3K9la peaks for *Sell, Bach2, Il7r*, and *Klf2* gene loci in naïve CD8 T cells treated with and without 25mM Na-La. Whiskers for all box-and-whisker plots in figure S8 represent minimum-maximum values.

**Figure S9:**
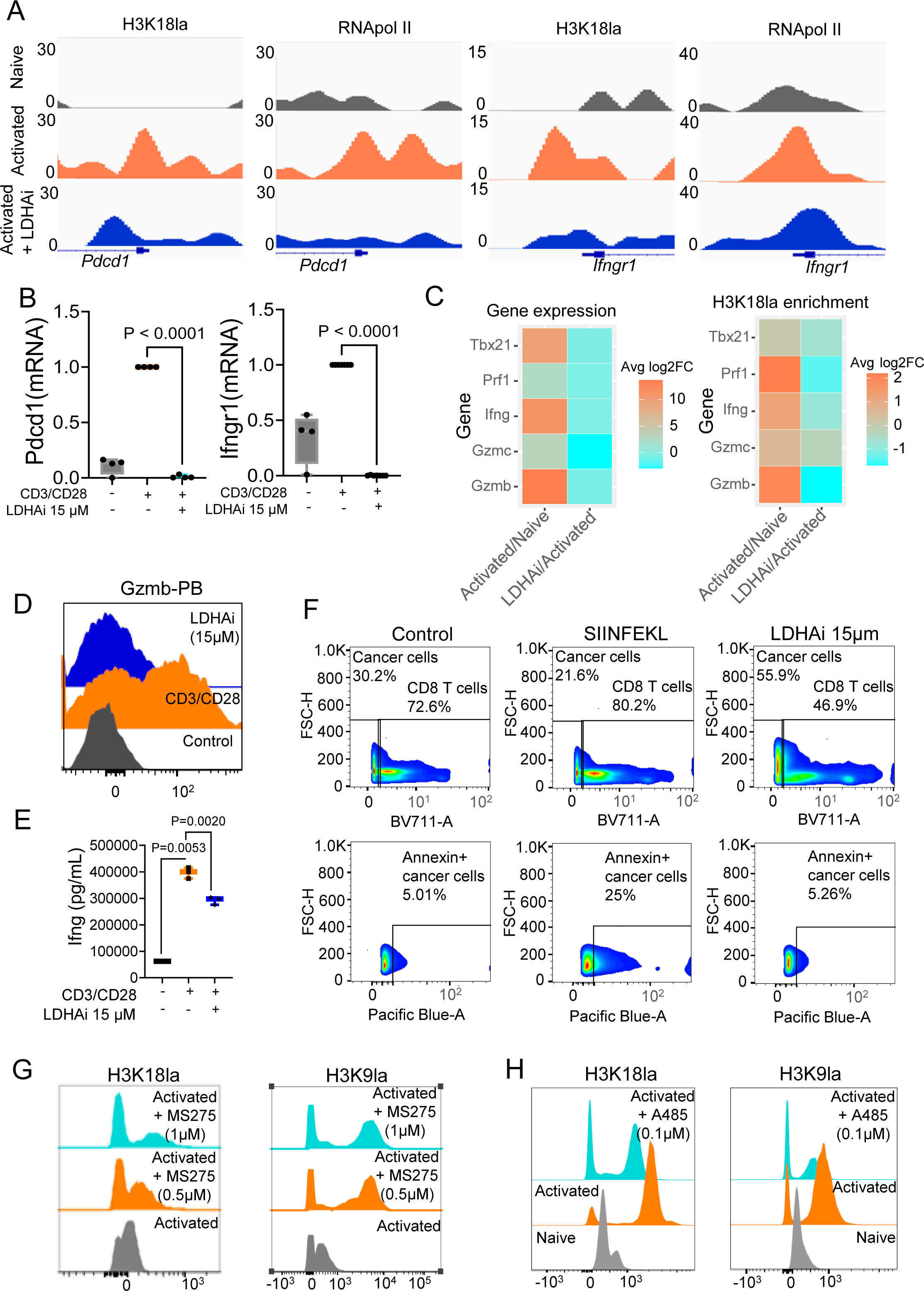
**A,** Genome Browser view comparing H3K18la and RNA pol II peaks at the TSS ±1kb regions of the indicated genes in naïve, activated and 15 µM LDHA treated (LDHAi) activated CD8 T cells. **B**, Box-and-whisker plots showing the relative expression of *Pdcd1* and *Ifngr1* in naïve, activated and LDHA inhibitor treated activated CD8 T cells normalized to the expression of β-actin as determined by quantitative PCR. (n=4) Two-tailed Student’s t-test was performed **C**, Heatmaps demonstrating the average log2Fold Change of gene expression (left panel), and H3K18la enrichment (right panel) following T cell activation (first column of both panels) and LDHA inhibitor treatment of activated CD8 T cells (second column of both panels). **D**, Representative histograms showing Granzyme B levels in naïve, activated and 15µM LDHA treated activated CD8 T cells. **E**, Box-and-whisker plot demonstrating the concentration of interferon gamma in culture supernatants of naïve, activated and 15 µM LDHA treated CD8 T cells. n=3 female mice. Two-tailed Student’s t-test was performed. **F**, (Top panel) Representative pseudocolor plots demonstrating the percentage of MB49-ova cancer cells and OT-I CD8 T cells following 24-hour co-culture of the cancer cells with unstimulated, SIINFEKL pre-stimulated and 15 µM LDHA inhibitor treated OT-I CD8 T cells. (Bottom panel) Representative pseudocolor plots depicting the percentage of annexin-V positive MB49-ova cancer cells following 24-hour co-culture with unstimulated, SIINFEKL pre-stimulated and 15 µM LDHA inhibitor treated OT-I CD8 T cells. **G,** Histograms showing MFI of H3K18la and H3K9la in activated, 0.5 µM and 1 µM MS275 treated activated CD8 T cells. **H,** Histograms depicting MFI of H3K18la and H3K9la in naïve, activated and 0.1 µM A485 treated activated CD8 T cells. Whiskers for all box-and-whisker plots in figure S9 represent minimum-maximum values.

**Supplementary Table 1:**
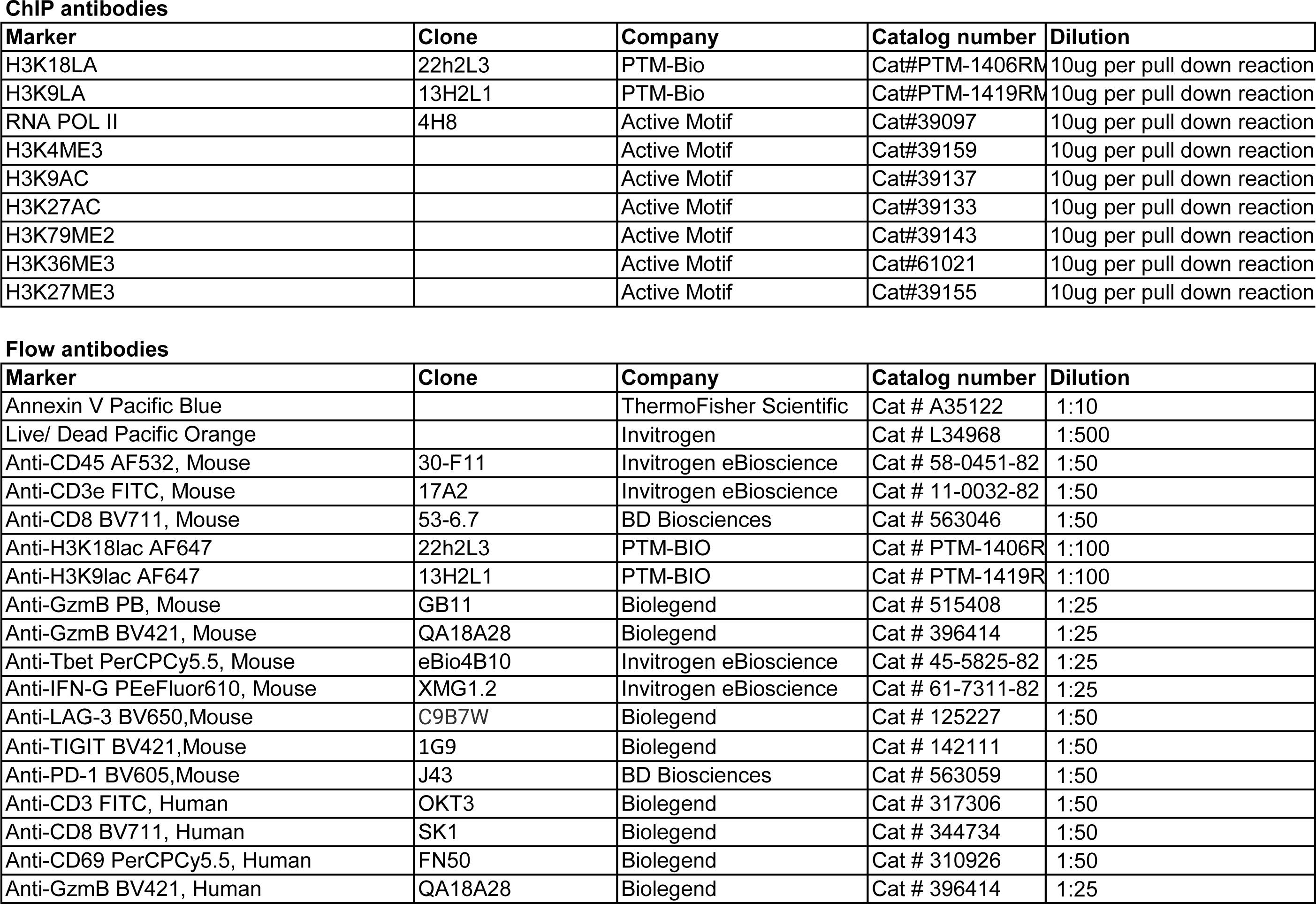
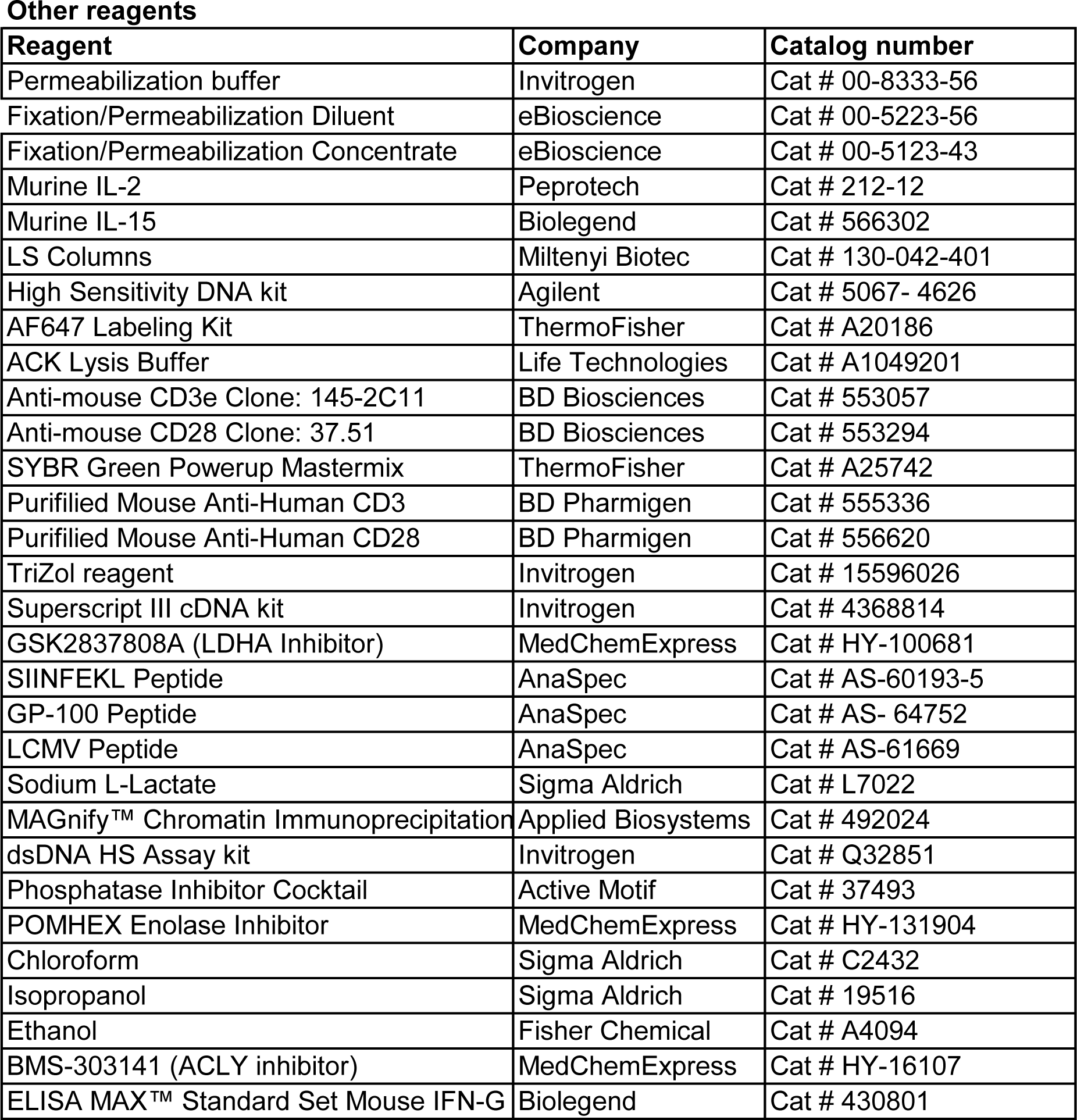
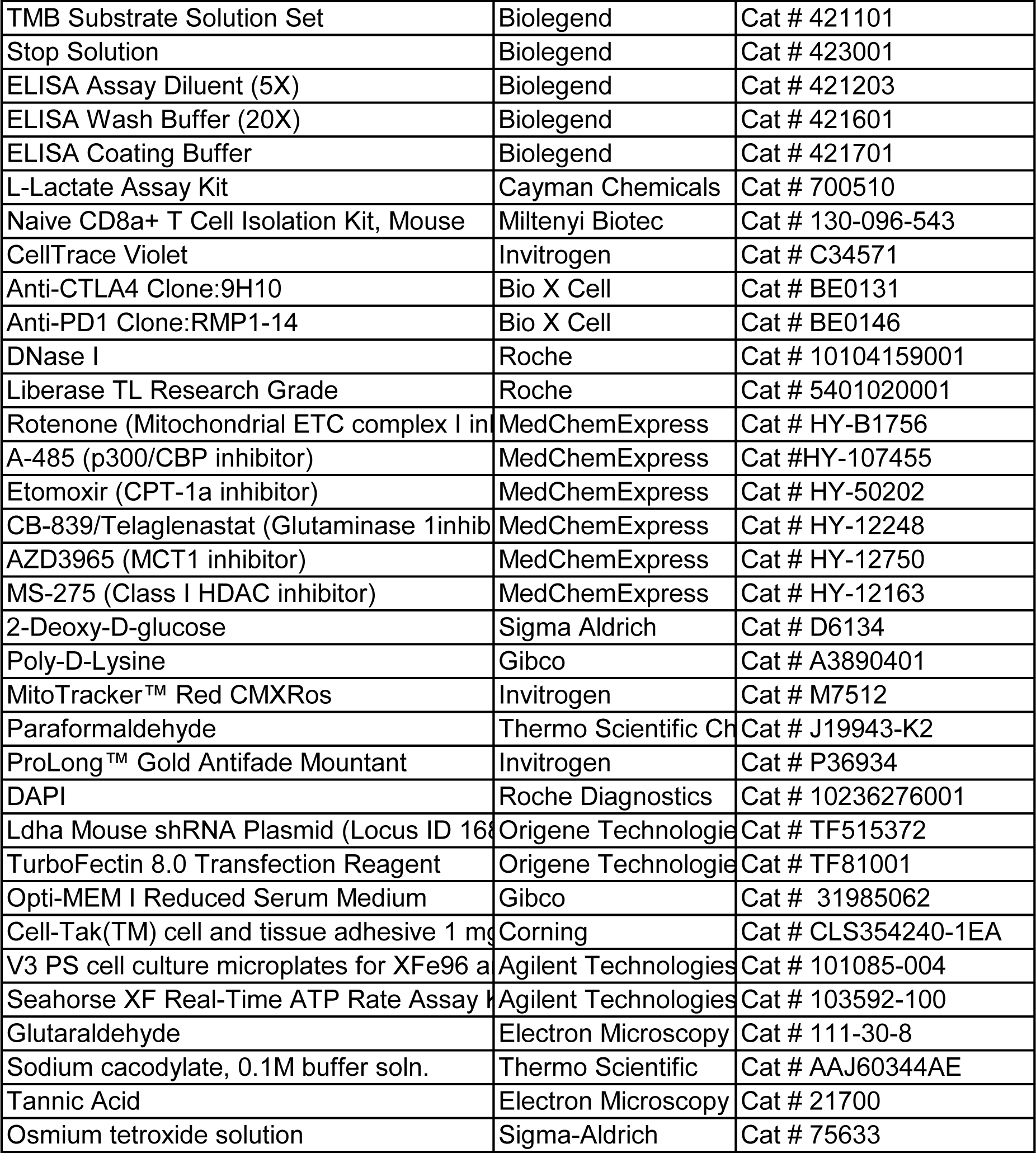
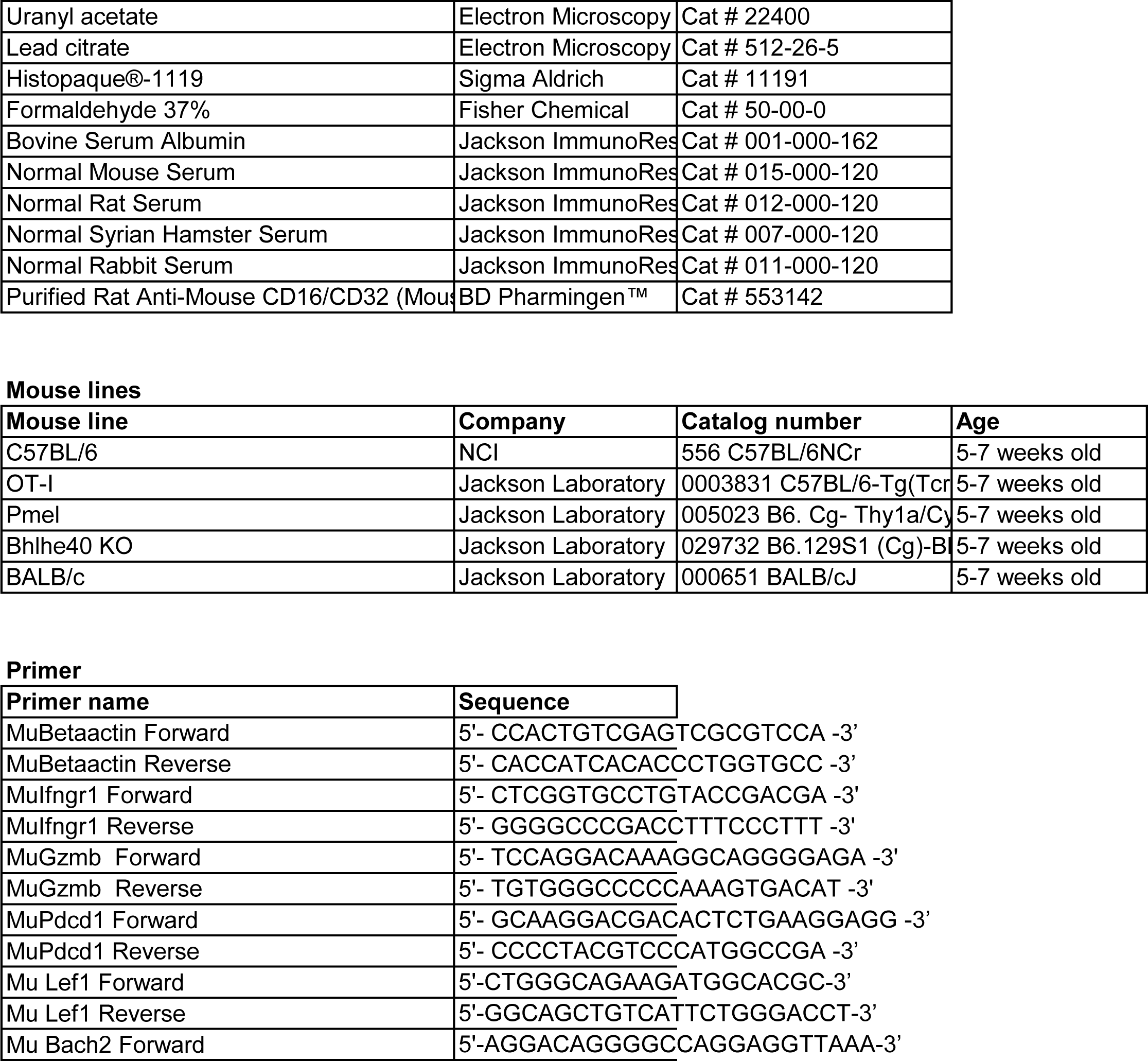
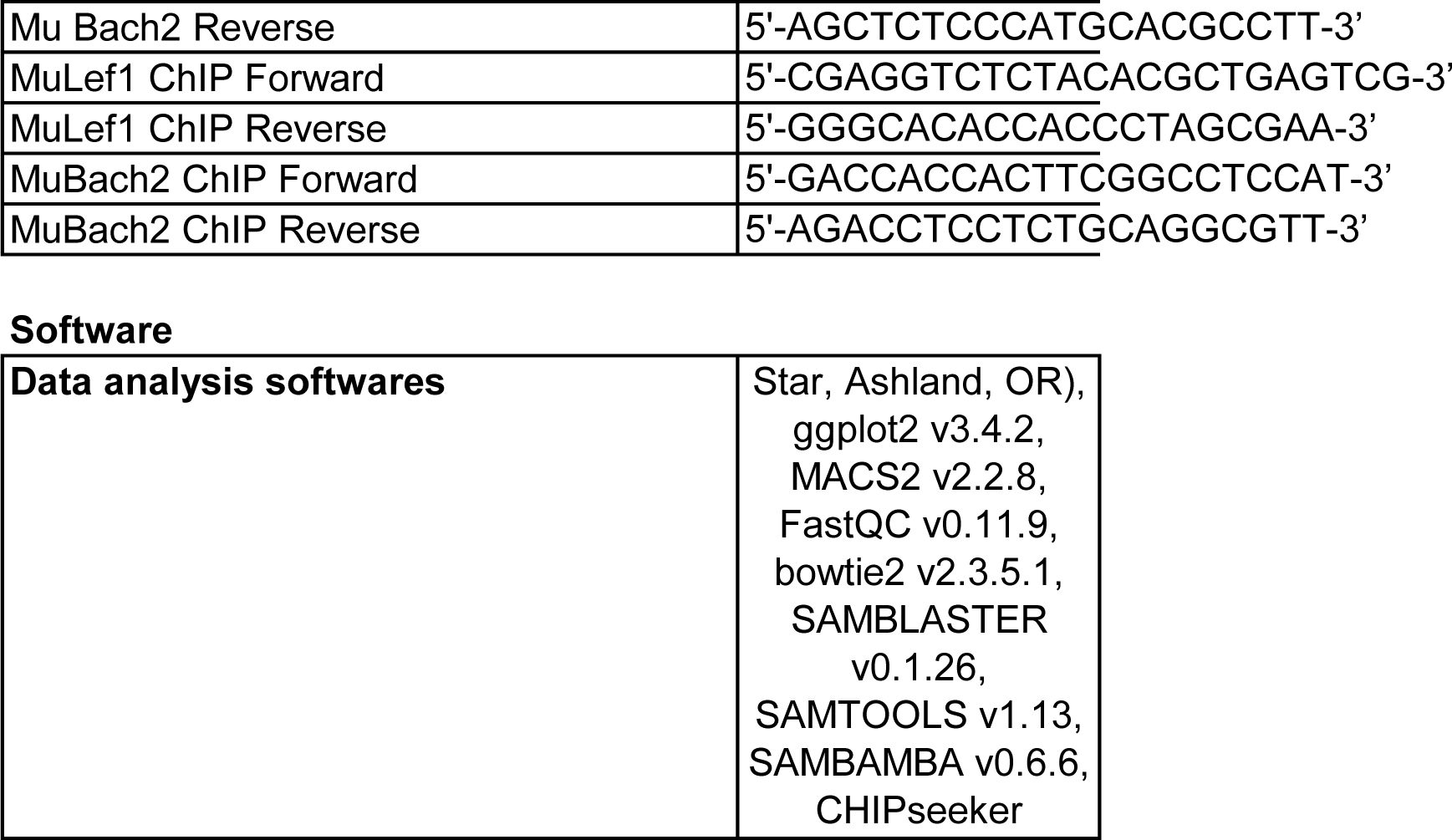
Key Resource table with list of antibodies.

